# Evolutionary divergence of the Wsp signal transduction system in β- and γ-proteobacteria

**DOI:** 10.1101/2021.07.02.450980

**Authors:** Collin Kessler, Eisha Mhatre, Vaughn Cooper, Wook Kim

## Abstract

Bacteria rapidly adapt to their environment by integrating external stimuli through diverse signal transduction systems. *Pseudomonas aeruginosa*, for example, senses surface-contact through the Wsp signal transduction system to trigger the production of cyclic di-GMP. Diverse mutations in *wsp* genes that manifest enhanced biofilm formation are frequently reported in clinical isolates of *P. aeruginosa*, and in biofilm studies of *Pseudomonas* spp. and *Burkholderia cenocepacia*. In contrast to the convergent phenotypes associated with comparable *wsp* mutations, we demonstrate that the Wsp system in *B. cenocepacia* does not impact intracellular cyclic di-GMP levels unlike that in *Pseudomonas* spp. Our current mechanistic understanding of the Wsp system is entirely based on the study of four *Pseudomonas* spp. and its phylogenetic distribution remains unknown. Here, we present the first broad phylogenetic analysis to date to show that the Wsp system originated in the β-proteobacteria then horizontally transferred to *Pseudomonas* spp., the sole member of the γ-proteobacteria. Alignment of 794 independent Wsp systems with reported mutations from the literature identified key amino acid residues that fall within and outside annotated functional domains. Specific residues that are highly conserved but uniquely modified in *B. cenocepacia* likely define mechanistic differences among Wsp systems. We also find the greatest sequence variation in the extracellular sensory domain of WspA, indicating potential adaptations to diverse external stimuli beyond surface-contact sensing. This study emphasizes the need to better understand the breadth of functional diversity of the Wsp system as a major regulator of bacterial adaptation beyond *B. cenocepacia* and select *Pseudomonas* spp.

**Importance:** The Wsp signal transduction system serves as an important model system for studying how bacteria adapt to living in densely structured communities known as biofilms. Biofilms frequently cause chronic infections and environmental fouling, and they are very difficult to eradicate. In *Pseudomonas aeruginosa*, the Wsp system senses contact with a surface, which in turn activates specific genes that promote biofilm formation. We demonstrate that the Wsp system in *Burkholderia cenocepacia* regulates biofilm formation uniquely from that in *Pseudomonas* species. Furthermore, a broad phylogenetic analysis reveals the presence of the Wsp system in diverse bacterial species, and sequence analyses of 794 independent systems suggest that the core signaling components function similarly but with key differences that may alter what or how they sense. This study shows that Wsp systems are highly conserved and more broadly distributed than previously thought, and their unique differences likely reflect adaptations to distinct environments.

## Introduction

Biofilms are extremely recalcitrant in nature, which is driven largely by the extracellular matrix comprising diverse compounds produced by individual cells^1–8^. This matrix manifests structured community growth and dynamically generates sharp chemical gradients to drive both phenotypic and genetic diversification. Exopolysaccharides (EPS) are a major component of this matrix, and they play a critical role in cell-cell and cell-surface adhesion^9–12^. EPS production or export is modulated by the second messenger cyclic di-GMP^5,13–21^ in many organisms, which promotes opportunistic pathogens like *Pseudomonas aeruginosa* to persist for years in the lungs of cystic fibrosis patients and cause extensive damage^18,22,23^. Clinical isolates of *P. aeruginosa* often display phenotypic heterogeneity as either smooth, mucoid, or small colony variant (SCV) phenotypes^22,24–26^. Sequence analyses of SCVs commonly identify mutations in *wsp* (<u>w</u>rinkly <u>s</u>preader <u>p</u>henotype) genes that increase the intracellular pool of cyclic di-GMP^27–31^.

Studies of the Wsp signal transduction system in several *Pseudomonas* species have played an important role in establishing the positive correlation between cyclic di-GMP production and biofilm formation^18,19,27,28,32–37^. The *Pseudomonas* Wsp system is encoded by the *wspABCDEFR* monocistronic operon that relays surface contact as an extracellular signal^38^ to activate the diguanylate cyclase activity of WspR to produce cyclic di-GMP, which in turn stimulates EPS production and biofilm formation^5,18–21^. Currently, the functional model of the Wsp system (Fig. 1A) is largely based on sequence similarity to the respective Che proteins that collectively make up the enteric chemotaxis system^28^. The trans-membrane component of the Wsp complex, WspA, forms a trimer-of-dimers that is laterally distributed around the cell and acts to initiate cyclic di-GMP production^34^. When activated by cellular contact with a surface, WspA is predicted to undergo a conformational change to expose its methylation sites, and a methyltransferase (WspC) and a methylesterase (WspF) ultimately determine the active state of WspA^19^. Methylated WspA initiates the autophosphorylation of the histidine kinase WspE^27,34,37^, which then phosphorylates the diguanlyate cyclase WspR to activate cyclic di-GMP production. WspE also phosphorylates WspF which terminates the signaling cascade^27,34,35^. The Wsp complex is predicted to be functionally stabilized by WspB and WspD which anchor the WspA trimer-of-dimer complex to WspE.

**Figure 1.**
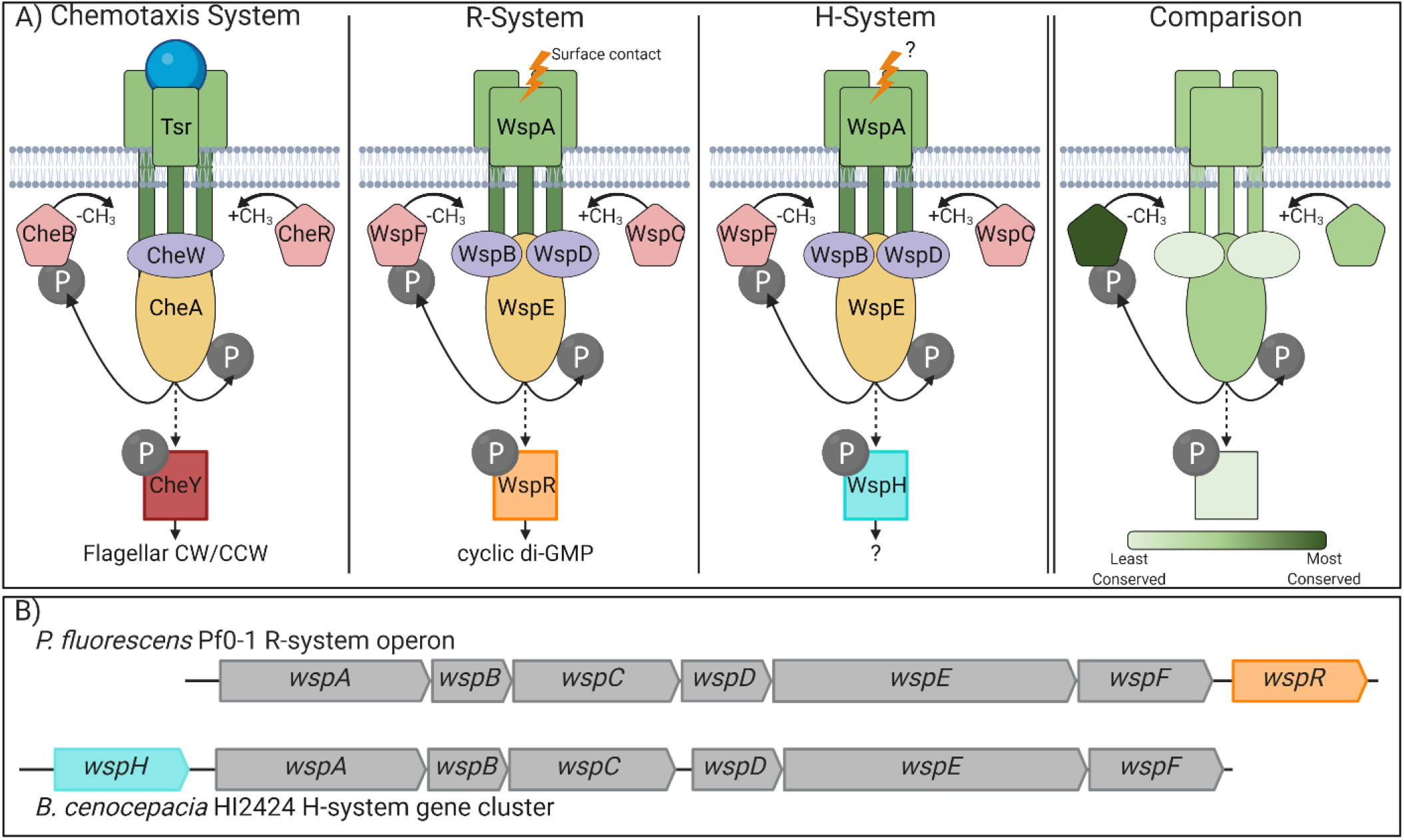
Comparison of the Wsp signal transduction system to the enteric chemotaxis system. A) A schematic comparison of the chemotaxis (Che) system of *E. coli* to the WspR system of *Pseudomonas* and the WspH system of *Burkholderia*. The Che system modulates the direction of flagellar rotation in response to the binding of attractants to the receptor (e.g. serine and Tsr). The Wsp system is reported to respond to surface contact in *P. aeruginosa*, but the signal output varies between activating WspR (diguanylate cyclase) or WspH (function unknown). The panel on the right depicts sequence conservation of relative proteins among the Che, WspR, and WspH systems as presented numerically in Table S1. Dark green proteins show the greatest conservation while light green proteins show the least conservation. CH3 = methyl group, P= phosphate. B) The *wsp* genes in *P. fluorescens* Pf0-1 and *B. cenocepacia* HI2424 share synteny except that *wspR* is the terminal gene of a monocistronic operon and *wspH* is absent in the latter; *wspH* appears to be encoded as an independent transcription unit upstream from the *wsp* operon.

**Figure 2.**
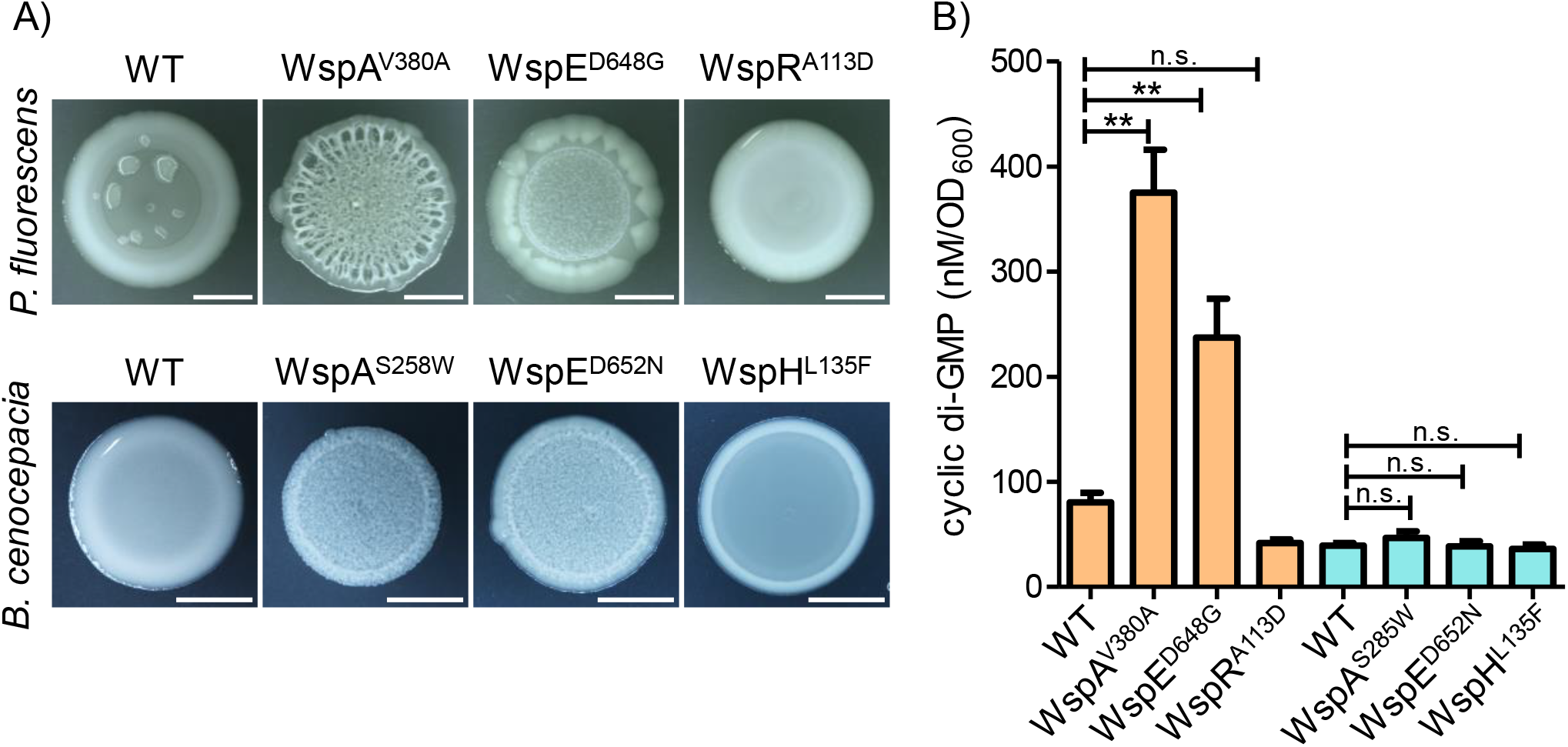
Comparable Wsp mutants of *P. fluorescens* and *B. cenocepacia* produce similar colony morphologies but differ in cyclic di-GMP production. A) Colony images of *P. fluorescens* and *B. cenocepacia* (day 4) show that mutations within the *wspA* or *wspE* genes in either organism produce the wrinkled phenotype while mutations in *wspH or wspR* produce a smooth phenotype (scale bar = 5mm). The mucoid patches observed in the WT *P. fluorescens* colony represent *rsmE* mutants that naturally emerge (Kim *et. al*, 2014). B) LC-MS/MS data show that the wrinkly phenotype correlates with increased levels of cyclic-di-GMP in Wsp mutants in *P. fluorescens* (orange bars) but not in *B. cenocepacia* (teal bars). Plotted are the mean of three replicates, error bars represent the standard deviation, n.s. denotes no significant difference, and ** denotes significant difference (ANOVA p < .0001; TukeyHSD p < 0.01).

Although multiple studies have empirically demonstrated overarching functional parallelism in signal transduction between the Wsp and enteric chemotaxis systems^19,27,28,34,36,39–42^, fundamental gaps remain in our mechanistic understanding. For example, WspB and WspD are assumed to be functionally redundant as they are both homologous to CheW. However, this appears not to be the case since the subcellular localization of WspA differs between Δ*wspB* and Δ*wspD* mutants^19^. Furthermore, all studies of the Wsp system had been limited to four *Pseudomonas* spp. until the discovery that *Burkholderia cenocepacia* HI2424 lacks the *wspR* gene and instead possesses *wspHRR* (hereafter, *wspH*), a locus predicted to encode a unique hybrid histidine kinase/response regulator without the diguanylate cyclase domain^43,44^. Despite lacking WspR, mutations in the *wsp* genes of *B. cenocepacia* HI2424 produce the wrinkly colony morphologies and the SCV phenotypes similar to the *wsp* mutations in *Pseudomonas* spp.^43,44^. Although these observations imply that the Wsp system in *B. cenocepacia* HI2424 also functions to regulate cyclic di-GMP production, its cognate diguanylate cyclase, if any, remains unknown.

Since the initial characterization of the Wsp system in *P. fluorescens*^27,28,32–34,45^, mutations within each of the *wsp* genes in other species have been reported to alter both the colony morphology and biofilm phenotypes, and they were either demonstrated or presumed to impact cyclic di-GMP production^19,37,39,46,47^. Here, we show for the first time that mutations within the *wsp* operon of *B. cenocepacia* HI2424 do not alter the intracellular pool of cyclic di-GMP despite producing similar wrinkly colony phenotypes as comparable mutants in *Pseudomonas* spp. The function of the Wsp system may have further diverged beyond *Pseudomonas* and *Burkholderia* spp., but this remains entirely unexplored. To gain a broader appreciation of the mechanistic and functional robustness of Wsp systems, we first construct a dataset of taxonomically diverse Wsp systems and assess their phylogenetic distribution and evolutionary history. We then evaluate sequence conservation at each amino acid residue to identify conserved genetic elements both within and outside annotated functional domains. By anchoring our sequence conservation analysis with diverse *wsp* mutations across multiple species that are known to impact signal transduction, we re-evaluate the current functional model of the Wsp system and identify specific amino acid residues that likely manifest the key mechanistic differences between cyclic di-GMP dependent and independent Wsp systems.

## Methods

### Strains and culture conditions

All bacterial strains were routinely grown in Lennox LB (Fisher BioReagents) or on Pseudomonas Agar F (PAF) (Difco). Pseudomonas F (PF) broth (a non-agar variant of PAF) was prepared with the following composition: pancreatic digest of casein 10 g/L (Remel), proteose peptone No. 3 10 g/L (Remel), dipotassium phosphate 1.5 g/L (Sigma-Aldrich), and magnesium sulfate 1.5 g/L (Sigma-Aldrich). *Pseudomonas* and *Burkholderia* strains were incubated at 30°C and 37°C, respectively, with 250 rpm shaking when applicable. *wsp* mutants used in this study were previously isolated from *Burkholderia cenocepacia* HI2424^43^ and *Pseudomonas fluorescens* Pf0-1^39^. Colony morphology was evaluated in triplicate by inoculating PAF plates (9 mm petri dish, 25 mL PAF) with 25 μl of overnight culture then incubating at 22°C for 4 days.

### Cyclic di-GMP extraction and LC-MS/MS

An isolated colony of each strain was transferred to PF broth and grown overnight at the respective temperature. Overnight cultures were diluted to an OD_600_ of 0.04 in triplicate and incubated at the respective temperature for 2-4 hours until the samples reached an OD_600_ of 0.5, then promptly processed for cyclic di-GMP extraction. Cyclic di-GMP extraction and quantification procedures followed the protocols established by the Michigan State University Research Technology Support Facility (MSU-RTSF): MSU_MSMC_009 protocol for di-nucleotide extractions and MSU_MSMC_009a for LC-MS/MS. All following steps of cyclic di-GMP extraction were carried out on ice and noted otherwise. 1mL of 0.5 OD_600_ culture was centrifuged at 15,000 RCF for 30 seconds, and cell pellets were resuspended in 100uL of an ice cold acetonitrile/methanol/water (40:40:20 (v/v/v)) extraction buffer supplemented with a final concentration of 0.01% formic acid and 25nM cyclic di-GMP-Fluorinated internal standard (InvivoGen CAT: 1334145-18-4). Pelleted cells were mechanically disturbed with the Qiagen TissueLyser LT at 50 oscillations/second for 2 minutes. Resuspended slurries were incubated at −20°C for 20 minutes and pelleted at 15,000 RCF for 15 minutes at 4°C. The supernatant was transferred to a pre-chilled tube then supplemented with 4uL of 15% w/v ammonium bicarbonate (Sigma-Aldrich) buffer for stable cryo-storage^47^. Extracts were stored at −80°C for less than two weeks prior to LC-MS/MS analysis at MSU-RTSF. We observed sample degradation with repeated freeze-thaw cycles, so we ensured that the extracts were never thawed prior to LC-MS/MS analysis. All samples were evaporated under vacuum (SpeedVac, no heat) and redissolved in 100uL of the mobile phase (10 mM TBA/15 mM acetic acid in water/methanol, 97:3 v/v, pH 5). UPLC/MS/MS quantification used Waters Xevo TQ-S instrument with the following settings: cyclic di-GMP-F at *m/z* transition of 693 to 346 with cone voltage of 108 and collision voltage of 33; cyclic di-GMP at *m/z* transition of 689 to 344 with cone voltage of 83 and collision voltage of 31. Cyclic di-GMP data was normalized to 25nM cyclic di-GMP-F by MSU-RTSF to account for sample matrix effects and sample loss during preparation. We used the OD_600_ measurements of the initial samples to report the quantified cyclic di-GMP as nM/OD and visualized with GraphPad Prism5.

### Generation of the H- and R-systems dataset

The WspH encoding system of *B. cenocepacia* HI2424 and the WspR encoding system of *P. fluorescens* Pf0-1 were used as queries to search the BCT bacterial subdivision of GenBank for syntenic homologs (NCBI protein accession numbers: [ABK10522.1, ABK10523.1, ABK10524.1, ABK10525.1, ABK10526.1, ABK10527.1, ABK10528.1, ABK10529.1] and [ABA72795.1, ABA72796.1, ABA72797.1, ABA72798.1, ABA72799.1, ABA72800.1, ABA72801.1], respectively). All previous Wsp studies show that the *wsp* operon is highly syntenic ^19,27,28,32–34,37,39,45–47^ and *B. cenocepacia* HI2424 shares this synteny^43^. Cooper *et al*. identified that *wspH* had relocated from the 3’ end to the 5’ end of the WspH gene cluster^43^. To determine if syntenic homologs have a similar *wspH* gene cluster arrangement and to generate a large dataset of H- and R-systems, we used MultiGeneBlast v-1.1.14 to search GenBank and identify homologous Wsp systems. MultiGeneBlast blasts for query sequences and then provides a weighted score based on the synteny of the genes^48^. We executed MultiGeneBlast locally against the bacterial subdivision of GenBank (BCT subdivision) and reported the first 2,000 syntenic homolog hits for the WspH encoding system of *B. cenocepacia* HI2424 and the WspR encoding system of *P. fluorescens* Pf0-1. The *P. fluorescens* Pf0-1 search results are provided as Data S1 and the *B. cenocepacia* HI2424 search results are provided as Data S2. Our search produced 1640 R-system hits and 1500 H-system hits. Both searches generated less than 2,000 hits as we had specified, indicating that we have identified nearly all available *wsp* homologs in the GenBank database that match our queries.

Assessment of the R-system dataset showed that the first 810 R-system hits possessed a GGDEF or GGEEF domain and shared synteny with the *P. fluorescens* Pf0-1 queries. Hits 811-812 of the R-system search identified species that did not contain a *wspR* response regulator hit because an entry for that locus was not included in their GenBank annotation files. Hits 813-827 and 829 identified *Burkholderia* spp. with a 5’ response regulator that lacked a GGDEF or GGEEF domain and closely resembled the *wspH* gene cluster of *B. cenocepacia* HI2424. Hits ≥830 did not contain a gene cluster with similarity to the *wspR* or *wspH* gene sets. Based on this information, we collected the first 810 R-system hits to be included in the final R-system dataset. A similar distribution was observed in the 1500 H-system hits. The first 217 H-system hits identified were *Burkholderia* species with a 5’ response regulator that lacked a GGDEF or GGEEF domain. Hits 218-818 included a GGDEF or GGEEF domain and closely resemble the *wspR* gene cluster of *P. fluorescens* Pf0-1. Hits 819 to 829 were *Pseudomonas* species that did not contain a *wspH* hit. Manual analysis identified these as *wspR* containing operons with low *wspH* homology. Hits ≥830 did not contain a gene cluster with similarity to the *wspR* or *wspH* gene sets. Based on this information, we collected the first 818 H-system hits to be included in our H-system dataset. Compiling the two datasets and removing duplicate hits generated a final dataset of 794 total hits. We identified 588 R-systems based on the presence of the GGDEF or GGEEF domain and 206 H-systems that lacked this domain and were 5’ adjacent. We did not identify any H-systems with a 3’ response regulator nor did we find any organisms that contained both H- and R-systems.

### Phylogenetic analysis of WspH/R dataset strains

The MultiGeneBlast output includes the NCBI organism accession number and the protein accession numbers for each sequence identified in the analysis. The RefSeq URL web address for the assembly statistics file of each organism was determined from the NCBI organism accession number using the Entrez.esearch and Entrez.read utility of BioPython v-1.78^49^ (code provided as File S1). Data for *Caulobacter crescentus* (Table S2) was obtained manually for rooting the species phylogeny. We chose *C. crescentus* because it is an α-proteobacterium that contains the diguanylate cyclase PleD which has been previously used as a reference in *wspR* studies^41^. R statistical software was then used to download all RefSeq assembly data files for each organism using generated URL addresses based on the assembly statistics files (code provided as Files S2, S3, and S4).

The obtained files were compared to the Bac120 ubiquitous housekeeping gene set established by Parks *et al*^50^. This search was conducted with the hmmsearch tool of HMMER v-3.3.2 with an e-value cutoff of 1e-10^51^. The HMM files were converted to sequence files using the esl.sfetch tool of the Easel HMMER library using default settings. Some organisms contained multiple copies of a housekeeping gene. In these instances, the gene with the lowest e-value was select for the analysis. Organisms that lacked any of the Bac120 genes received a blank fasta entry indicated by dashes. Peptide encoding sequences for each gene in the Bac120 set were aligned using MAFFT v-7.475 with 1000 iterations and global pair options^52^. Trimal v-1.3 was then used to remove gaps with a 0.9 cutoff value option^53^. This generated 120 individual alignments with 795 organisms per alignment (794 dataset plus rooting sequence). Finally, one continuous sequence was generated for each organism by concatenating the 120 aligned sequences into a single string. This ultimately generated a single file with 795 organisms where one entry included the aligned information of all 120 encoded genes. This method has much greater sensitivity to differentiate species than the traditional 16S phylogenetic methods^50^. The final alignment was evaluated under the maximum likelihood (ML) algorithm RAxMLv-8.0.0 (HPC-HYBRID-AVX2) with the PROTCATBLOSUM62 model, 100 bootstraps, and the -f a hill-climbing options enabled. Final trees were rendered using the ITOL software suite (Fig. 3).

**Figure 3.**
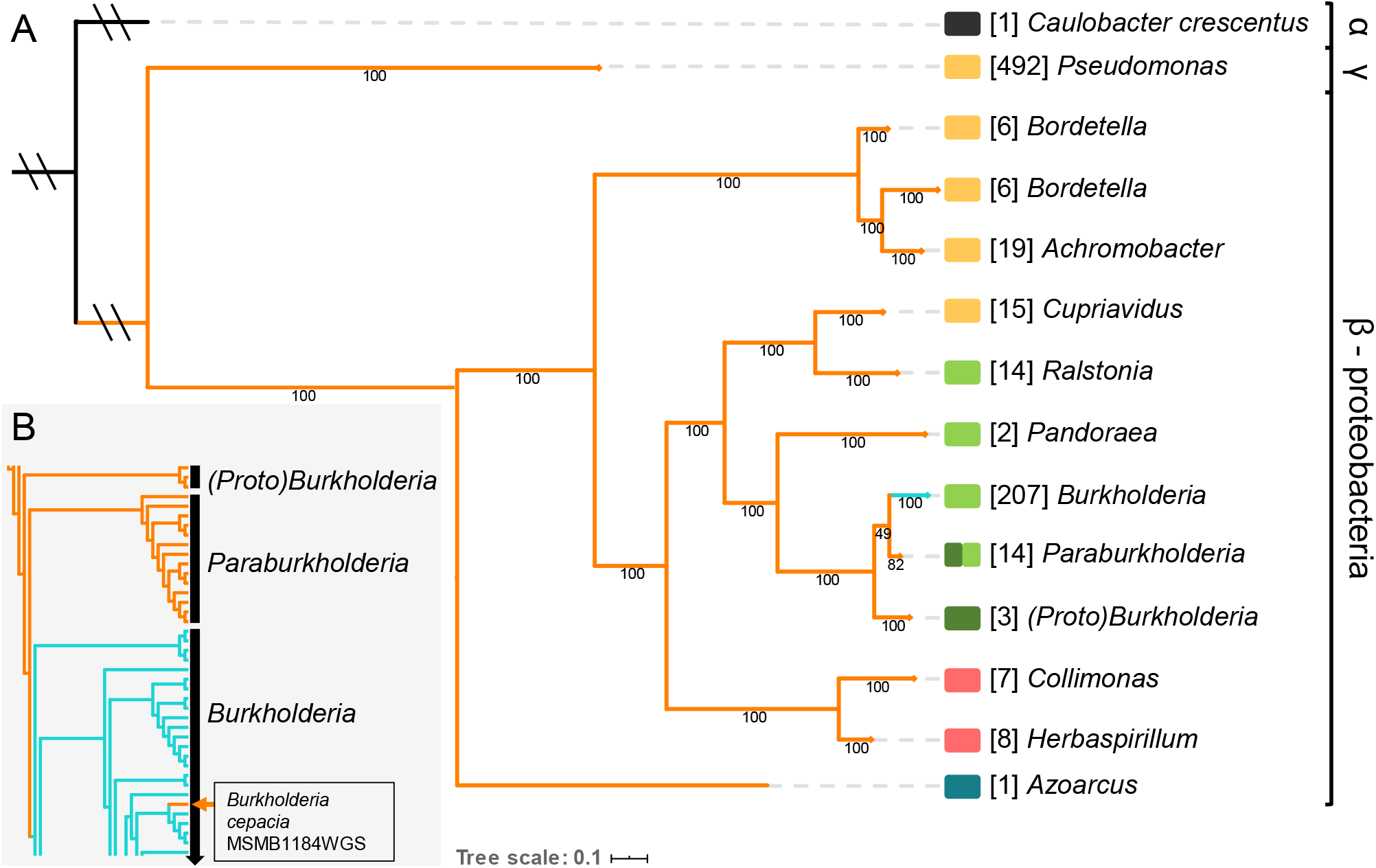
Phylogenetic analysis shows that the Wsp system is restricted to the β- and γ-proteobacteria and *wspH* is unique to *Burkholderia*. A) A maximum likelihood species tree of organisms with the H-system (teal) or the R-system (orange). For simplicity, the tree was collapsed at the genus level with the values within parentheses indicating the number of strains in each branch (detailed in Table S3). 588 strains possess the R-system, and 206 strains possess the H-system which is restricted to *Burkholderia*. B) An expansion of the *Burkholderia-Paraburkholderia* subgroup shows that only one strain (*Burkholeria cepacia* MSMB1184WGS) possesses the R-system. An independent phylogenetic assessment of Wsp proteins identified five unique Wsp system clades (Fig. S1): blue = clade 1, yellow = clade 2, pink = clade 3, light green = clade 4, dark green = clade 5. Values reported under individual branches are bootstrap support values (out of 100).

### Phylogenetic analysis of the Wsp signaling core

MultiGeneBlast provides protein accession numbers for sequences identified in the analysis. Sequence files for these peptides were downloaded from NCBI with the provided accession numbers using the Entrez.esearch, Entrez.read, and Entrez.fetch utilities of BioPython^49^ (code provided as File S1). As in the species tree, *C. crescentus* was selected to root the tree. *P. fluorescens* Pf0-1 *wsp* homologs were identified in *C. crescentus* and are described in Table S2. Fasta files were independently generated for WspA, WspB, WspC, WspD, WspE, or WspF, resulting in 795 sequences per file (794 dataset plus rooting sequence). WspR and WspH were omitted from this analysis since they are divergent in both function and sequence. The fasta files were aligned using MAFFT with 1000 iterations and global pair options^52^. Trimal was then used to remove gaps with a 0.3 cutoff value option^53^. A continuous ‘Wsp signaling core’ sequence was generated for each organism by concatenating the six independently aligned Wsp sequences into a single string. RAxML (HPC-HYBRID-AVX2) was called with the PROTCATBLOSUM62 model, 1000 bootstraps, and the -f a hill-climbing options enabled. Final trees were rendered using the ITOL software suite (Fig. S1).

### Assessment of pathogens/opportunists in dataset

The 794 organisms in our dataset constitute 132 species as reported in Table S3. A literature review at the species level identified that 47 of the 132 species in our dataset can be classified as an opportunistic pathogen (58% of the total dataset). Common sites of human infection for each opportunistic pathogen were obtained from the literature and are reported in Table S3. The 47 species represent 7 genera (of 11 genera in dataset) that contain pathogenic species. This data is reported in Fig. S2 where the 7 genera represent x-axis labels and the percent of pathogenic and nonpathogenic species within each genus are shown. Common sites of infection for each pathogen are shown as a percent of the pathogenic species subset.

### Assessment of sequence conservation using Shannon entropy

To evaluate the conservation of residues in each Wsp protein, alignments were generated as described for the Wsp phylogeny except that *C. crescentus* was not included in the alignments. Generated alignments were uploaded to the Protein Residue Conservation Prediction tool established by Capra *et al* to calculate the Shannon entropy of each position within the alignment^54^. We used the Shannon entropy scoring method with a window size of 3, sequence weighting enabled, and BLOSUM62 matrix options. This tool evaluates the entropy in an alignment where sites with high variation have high Shannon entropy. The tool then scales the entropy to a range of (0,1) and subtracts this score from 1 so that higher scores indicate a greater conservation. This value is referred to as the conservation score. The conservation scores provided by this analysis were then downloaded and interpreted via Python and plotted via the matplotlib library for each Wsp alignment. Residues with a high conservative threshold (≥ 0.80) conservation score are highlighted in red. Available SNP data and annotation data were mapped to this graph to provide context on the function of these identified regions. We accomplished this by generating a consensus sequence of each alignment using the SeqIO library of BioPython v-1.78^49^ (code provided as File S1). We then individually mapped all available mutation data in the literature to the consensus sequences in a case by case basis using MEGA alignment software^55^ (Table S4). This provided a residue position of where mutations would fall in the alignment. Likewise, the consensus sequences were annotated using available Wsp literature^19,27,28,32– 34,37,39,45–47^, NCBI’s Conserved Domain Database (CDD)^56^, and through homology to the enteric chemotaxis Che system as reported in Table S5.

### Identifying residues essential for divergent H-system signaling

Alignments were constructed for the seven Wsp proteins (WspA, WspB, WspC, WspD, WspE, WspF, and WspH/R) using the 794 peptide sequences as previously detailed. The alignment for each Wsp protein was parsed via Python scripts into an H- or R-system alignment subset, resulting in 206 or 588 peptides per alignment, respectively. The parsing method retained the organization of the original 794 sequence alignment, allowing the H- and R-system alignment subsets to be directly compared for each Wsp protein. We generated a consensus sequence for all 14 alignments (7 Wsp proteins parsed into H- or R-system subsets) using the consensus algorithm of the AlignIO library in BioPython^49^. Next, we calculated the conservation scores of the alignments as previously described. We used the generated consensus sequence and the conservation score to create residue pairs for each position in the alignment, creating 2,899 residue pairs. For example, residue 360 of WspA has a serine in the H-system (conservation score = 0.99999) but a lysine in the R-system (conservation score = 0.80182). We identified residues that had a conservation score ≥ 0.80 in both the H- and R-system datasets which indicates that the residue is likely functionally important in both systems. This reduced the dataset to 971 residue pairs. We next considered mismatched residue pairs where a mutation likely occurred within the H-system during or after the evolutionary divergence of the H- and R-systems, further reducing the dataset to 178 residue pairs. 56 ambiguous residue pairs, or residues assigned ‘X’ in the consensus sequence, were omitted from our analysis, resulting in 122 residues pairs. We used the dataset of Pechmann *et. al* to identify conservative or non-conservative mutations in the H-system subset compared to the R-system subset^57^. 73 residue pairs constituted a conservative mutation from the R-system to the H-system and were filtered out of our dataset. The final dataset contained 43 residue pairs where each residue is highly conserved in its respective alignment subset; the residue pair identifies a non-conservative mutation within the H-system compared to the R-system that may be essential in cyclic di-GMP independent H-system signaling.

### Motif assessment of WspB and WspD

Assessment of conservation scores in WspB and WspD found that the conserved sites of these proteins were not similar. To visualize this difference, we collected the conservation scores of residues with a high conservative threshold (≥ 0.80) that were continuous for 3 or more residues. The identified islands of conservation plus 2 flanking residues on both ends were selected from the alignment to generate a sequence logo. Sequence logos were generated using WebLogo v-2.8.2^58^ with default settings and the logo range values for the sequences of interest. Comparisons of the motif sequence logos of WspB and WspD were conducted manually.

## Results and Discussion

### The WspR and WspH systems are functionally divergent

The core signaling components of the enteric chemotaxis (Che) system and the Wsp system in *Pseudomonas* spp. and *B. cenocepacia* are expected to be mechanistically similarly due to the high sequence conservation of the functional domains (Fig. 1A). However, the overall sequence homology among these three systems is relatively low, and the Wsp proteins between *P. fluorescens* Pf0-1 and *B. cenocepacia* HI2424 share greater homology (Table S1). This suggests that the Wsp system in *B. cenocepacia* HI2424 functions similarly to the *Pseudomonas* Wsp system, but the former lacks WspR and is instead predicted to phosphorylate WspH, which lacks WspR’s GGDEF enzymatic domain required for cyclic di-GMP production^43,44^ (Fig. 1). To test if the Wsp system in *B. cenocepacia* HI2424 regulates cyclic di-GMP production and/or activates an unidentified cognate diguanylate cyclase, we collected comparable *wsp* mutants from *P. fluorescens* Pf0-1 and *B. cenocepacia* HI2424 that are known to impact signal transduction. The selected strains possess *wspA* mutations in the WspA/B/D/E signaling domain, *wspE* mutations in the receiver (REC) domain at the phosphoacceptor site, or *wspH/R* mutations in their respective REC domains. The mutations in *wspA* (V380A) and *wspE* (D648G) are known to increase cyclic di-GMP production and biofilm formation in *Pseudomonas* spp.^19,32,39^. *B. cenocepacia* HI2424 mutants utilized here, *wspA* (S258W) and *wspE* (D652N), were previously isolated and demonstrated to increase biofilm formation^43,44^. Conversely, the mutation in *wspR* (A113D) is predicted to terminate Wsp signaling as the mutated protein could no longer be phosphorylated^35^. The mutation in *wspH* (L135F) has been shown to produce a smooth colony phenotype with defective biofilm formation^43^, suggesting a reduced signaling output, if any at all.

We first assessed colony morphologies of the *wsp* mutants for the iconic wrinkled phenotype that reflects increased cyclic di-GMP production across diverse species ^19,27,28,32–34,37,39,45–47^. Mutations in the *wspA* and *wspE* genes of both *P. fluorescens* Pf0-1 and *B. cenocepacia* HI2424 exhibit similar wrinkled phenotypes, while the *wspR* mutant of *P. fluorescens* Pf0-1 and the *wspH* mutant of *B. cenocepacia* HI2424 both exhibit smooth morphologies (Fig. 2A) as expected at low cyclic di-GMP levels. We next quantified the intracellular levels of cyclic di-GMP in the same set of mutants to test whether the altered colony morphologies indeed reflect increased cyclic di-GMP production. As predicted, cyclic di-GMP levels in *wspA* and *wspE* mutants of *P. fluorescens* Pf0-1 are significantly greater than the WT, unlike the *wspR* mutant (Fig. 2B). In contrast, none of the mutants of *B. cenocepacia* HI2424 significantly differ in cyclic di-GMP levels compared to the WT (Fig. 2B). This indicates that either the Wsp system in *B. cenocepacia* HI2424 does not regulate cyclic di-GMP production or that its influence on cyclic di-GMP production is rapidly buffered. Regardless, there is a clear contrast here between the colony morphologies and cyclic di-GMP levels in the two species. Although the wrinkled colony morphology was positively correlated with biofilm production in these *B. cenocepacia* HI2424 *wsp* mutants^44^, this appears to be achieved through a cyclic di-GMP independent mechanism. This is not particularly surprising given that WspH lacks a diguanylate cyclase domain and instead possesses a hybrid histidine kinase domain. A recent *wsp* phenotype suppressor study in *B. cenocepacia* HI2424 implicated that Bcen2424_1436 (RowR) is critical for WspH signaling and subsequent polysaccharide synthesis^44^. RowR is a predicted DNA-binding response regulator of an uncharacterized two-component transduction system, however its direct interaction with WspH remains to be resolved.

Despite the overall sequence similarity, the Wsp system in *B. cenocepacia* HI2424 appears functionally divergent from the WspR system in *Pseudomonas*. However, comparable *wsp* mutations in *B. cenocepacia* HI2424 and clinically persistent *Pseudomonas* spp. converge on biofilm formation, producing similar clinically relevant phenotypes^22,59,60^. To date, the WspH system has only been reported in *B. cenocepacia* HI2424^43,44^ and its relative uniqueness remains unknown. Similarly, the shared characteristic synteny of the *wsp* operon has only been described in four *Pseudomonas* strains^27,28,33,61^, and the extent of its taxonomic distribution also remains unknown.

### Wsp systems are exclusive to the β- and γ-proteobacteria and the WspH system is restricted to *Burkholderia*

To assess the phylogenetic distribution and the evolutionary history of the Wsp system, we constructed a database of Wsp homologs from all publicly available bacterial genomes in GenBank (see Methods). Sequence similarity alone makes it difficult to bioinformatically distinguish Wsp homologs from those that function in chemotaxis (Fig. 1A)^62,63^. Fortunately, chemotaxis genes are infrequently encoded as a single operon^64,65^ in contrast to all annotated *wsp* genes (Fig. 1B). We thus identified syntenic Wsp homologs of *P. fluorescens* Pf0-1 and *B. cenocepacia* HI2424 with at least 30% sequence identity. Our analysis discovered 794 unique *wsp* gene clusters with conserved *wspA-wspF* synteny. Importantly, using the *wsp* genes from *P. fluorescens* Pf0-1 (Data S1) or *B. cenocepacia* HI2424 (Data S2) as independent queries generated overlapping results, indicating that synteny is a robust search parameter for identifying previously unannotated Wsp systems. All 794 identified *wsp* gene clusters were associated with either a *wspR* homolog (588) or a *wspH* homolog (206) as observed in *P. fluorescens* Pf0-1 or *B. cenocepacia* HI2424, respectively (Fig. 1B). We also found that no identified genome contains both H- and R-systems, and either system is present at a single instance per genome. For the sake of simplicity, we refer to *wsp* clusters with a *wspH* homolog as H-systems and those with a *wspR* homolog as R-systems.

We next constructed a phylogenetic tree to visualize the taxonomic distribution of the H- and R-systems, which reveals that they are present exclusively in β- and γ-proteobacteria (Fig. 3A). The R-system is distributed across both the β- and γ-proteobacteria while the H-system is limited to *Burkholderia*. The twelve *Bordetella* strains in our dataset form two distinct clades, and three *Burkholderia* strains (i.e. Proto-*Burkholderia*) branch out earlier from the remaining *Burkholderia* and *Paraburkholderia* clades. Proto-*Burkholderia* possesses the R-system as do the *Paraburkholderia*, suggesting that the R-system pre-dates the H-system found exclusively in *Burkholderia*. Further expanding the *Burkholderia* clade shows that the R-system is present in only one strain (Fig. 3B), *B. cepacia* MSMB1184WGS, a soil isolate from the Australian Northern Territory. *Pseudomonas* is the lone genus representing the γ-proteobacteria, indicating that it likely acquired the R-system through horizontal gene transfer.

### Phylogenetic incongruence reflects multiple horizontal transfer events of the R-system

We evaluated potential horizontal transfer events of the Wsp system by assessing the incongruence between the species and Wsp phylogenies^66,67^. A phylogeny of the 794 Wsp systems was constructed using the concatenated peptide sequences of the core Wsp signaling proteins (WspA, WspB, WspC, WspD, WspE, and WspF), which diverges into five distinct clades (Table S2, Fig. S1). The sequence variations in the Wsp signaling core alone clearly differentiate the H- and R-systems even in the absence of WspH and WspR. Significant differences between the species (Fig. 3) and Wsp (Fig. S1) trees strongly suggest that the Wsp system has been subjected to multiple horizontal transfer events as summarized in Fig. 4.

**Figure 4.**
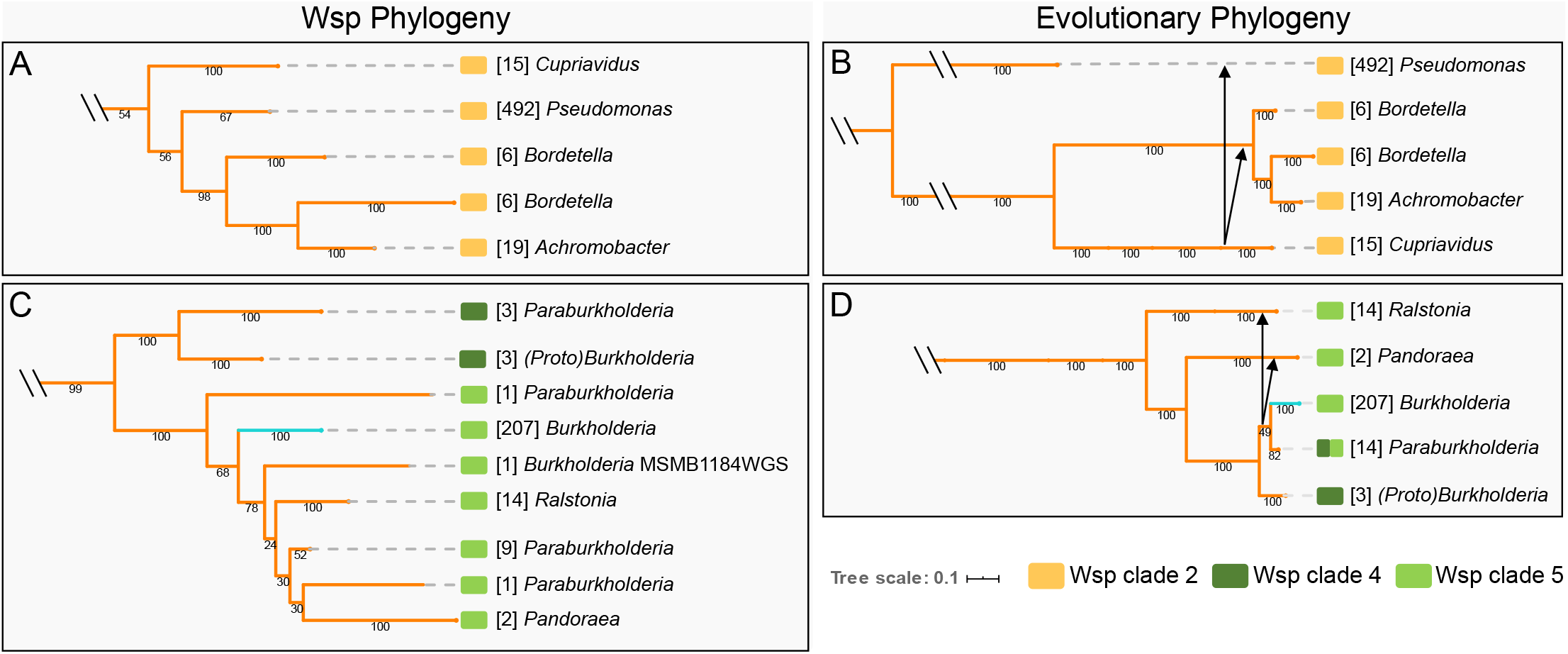
Phylogenetic analysis of the core Wsp proteins indicates multiple horizontal gene transfer events and establishes the R-system as the predecessor to the H-system. Shown here is a trimmed version of the Wsp phylogeny (A,C) for Wsp clades 2, 4, and 5 adjacent to the evolutionary phylogeny (B,D). The H-system (teal) and R-system (orange) Wsp phylogeny was constructed using the amino sequences of the core Wsp proteins (WspA, WspB, WspC, WspD, WspE, and WspF) and is detailed in Fig. S1. Clades identified in Fig. S1 have been reported as color blocks: yellow = Wsp clade 2, dark green = Wsp clade 4, and light green = Wsp clade 5. Values reported under branches are bootstrap support values (out of 100) and those in parentheses represent the total number of genomes/Wsp systems.

We observe that the R-system likely originated in *Azoarcus* (clade 1) then radiated throughout the β-proteobacteria and into *Pseudomonas* (Fig. S1), the sole member of the γ-proteobacteria (Fig. 3). In clade 2, the R-systems of *Pseudomonas, Bordetella*, and *Achromobacter* share a common node with *Cupriavidus* (Fig. 4A), yet these genera are taxonomically distinct (Fig. 4B). This observation, as supported by high bootstrap values in both phylogenies, indicates that *Cupriavidus* or its ancestor served as the common source for the horizontal transfer of the R-system into the phylogenetically distant *Pseudomonas, Bordetella*, and *Achromobacter* genera (Fig. 4B). Interestingly, each genus in clade 2 comprises opportunistic pathogens that are most frequently associated with the respiratory environment (Table S3, Fig. S2). The R-systems in clades 4 and 5 of the remaining β-proteobacteria appear to share a common ancestry (Fig. 4C) but show that the R-system from the ancestor of *Burkholderia* and *Paraburkholderia* likely transferred horizontally into *Ralstonia* and *Pandoraea* (Fig. 4D). This is clearly observed in the phylogenies where *Ralstonia* and *Pandoraea* taxonomically diverged before *Burkholderia* (Fig. 4D), but the acquisition of their Wsp systems occurred after *Burkholderia* (Fig. 4C). Interestingly, the Wsp system of *Paraburkholderia* is not confined to a single node in the Wsp phylogeny and is instead scattered among *Ralstonia, Pandoraea*, and *Burkholderia* (Fig. 4C). This indicates that the Wsp systems of *Paraburkholderia* have high sequence variation which could manifest unique functions such as responding to diverse stimuli or interacting with other response regulators.

Much like the taxonomic phylogeny, the H-system in *Burkholderia* forms a monophyletic clade, indicating that the signaling core of the H-system is highly conserved but is distinct from the R-system. The unique presence of the R-system in *B. cepacia* MSMB1184WGS presents an interesting case. Our analysis suggests that the H-system of this strain was independently replaced with an R-system after the radiation of the H-system in *Burkholderia* (Fig. 4C). Another possibility is that the R-system of this strain is related to the original source for the evolution of the H-system. Although many of the organisms in our dataset are associated with respiratory infections, we found that the Wsp systems are present in both opportunistic and non-pathogenic species (Table S3, Fig. S2). This suggests that horizontal transfer events of Wsp predate adaptations to the human host and that individual Wsp systems have functionally diverged beyond the emergence and integration of the *wspH* gene.

### Sequence conservation of Wsp systems and acquired mutations converge on predicted functional domains and unannotated regions

Missense mutations in *wsp* genes that impact biofilm formation are commonly identified in clinical isolates and experimental evolution studies^19,27,28,32–34,37,39,45–47^. Such mutations likely occur in key functional residues but the Wsp system as a whole remains poorly annotated. We thus aligned the amino acid sequence of all Wsp proteins in our dataset to generate a consensus sequence, and annotated functional domains based on homology to the enteric chemotaxis (Che) system and empirical studies of the *Pseudomonas* Wsp system (Tables S4 and S5). We then assessed the 794 sequence alignments for conservation using a weighted Shannon entropy algorithm where a conservation score near one indicates high conservation and a conservation score near zero indicates weak conservation^54^. Regions exceeding a conservation score of 0.8 are highlighted in red since this threshold reliably captures known functional residues^54^. Lastly, we compiled all *wsp* missense mutations from the literature that have been predicted to impact the signaling of the respective Wsp system, then mapped them to our consensus sequence (Table S4). We summarize the sequence conservation, newly annotated functional domains, and sites of missense mutations in Fig. 5, and the sequence conservation profiles of H- and R-systems independently in Fig. S3. The H-systems share consistently higher sequence conservation across all Wsp proteins, which reflects its phylogenetic isolation (Fig. S3). In general, our results complement previous reports and speculations on Wsp function but also reveal surprising patterns.

**Figure 5.**
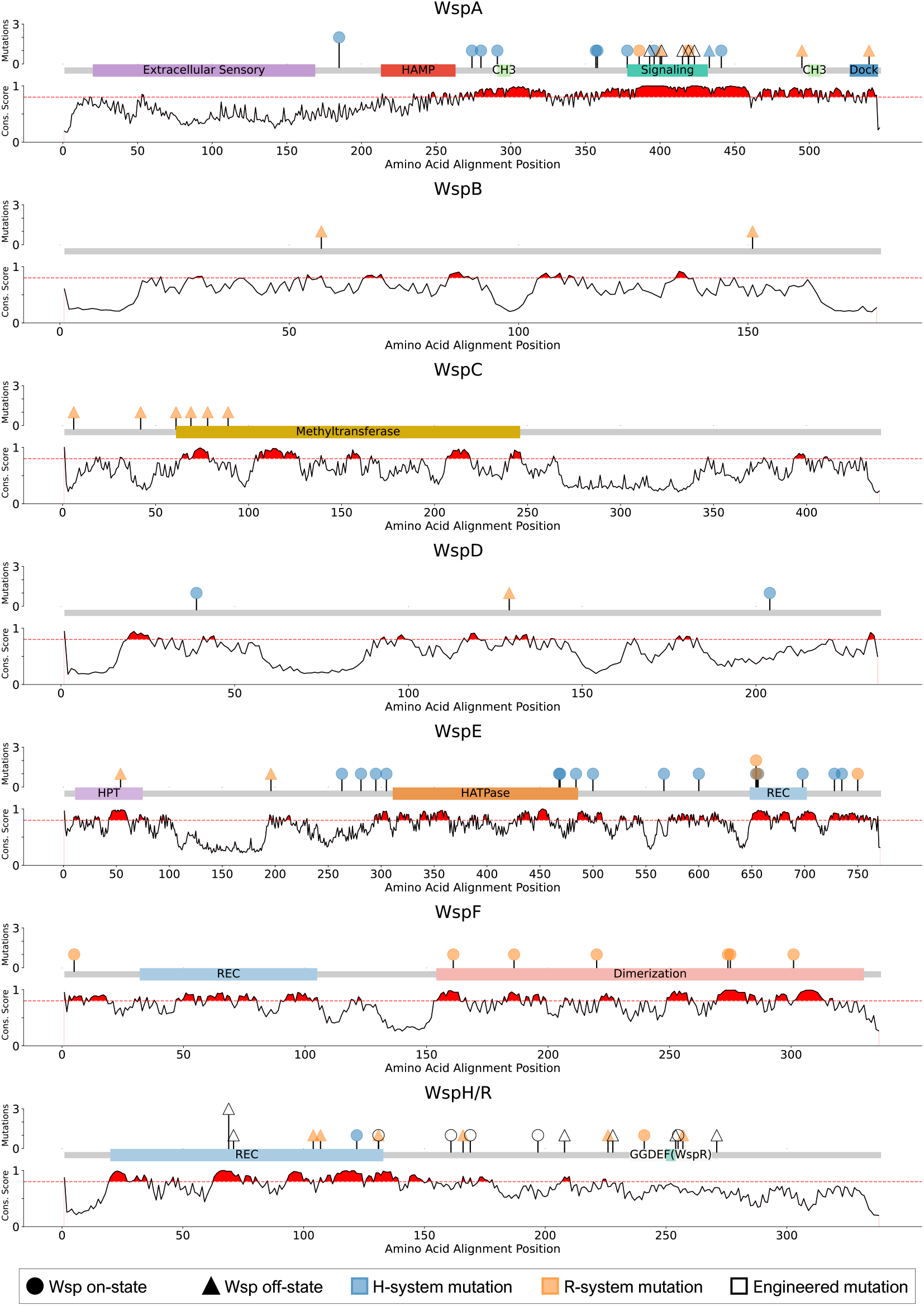
Evaluating the conservation of Wsp proteins identifies key residues that are likely essential to all Wsp signaling systems. The 794 peptide sequences for each protein were used to generate individual alignments. Annotation data is derived from NCBI CDD (conserved domain database), Prosite, or Che homology as indicated in Table S5. Reported naturally occurring missense mutations from the literature in the H-system are shown in blue and those in the R-system are shown in orange. Engineered missense mutations reported in the literature are indicated in white. Mutations that turn on the respective Wsp system are indicated as circles while those that turn off the system are indicated as triangles. The y-axis represents the Shannon Entropy evaluation for each protein alignment where weighted values near 1 indicate high sequence conservation and values near zero indicate weak sequence conservation. Regions where the weighted Shannon Entropy metric equals or exceeds 0.8 are shaded in red and denote regions likely to have functional or structural importance. Many of the previously identified mutations reside in these regions of high conservation but the functional role of these residues remains unknown.

WspA exhibits strikingly high sequence conservation across four distinct regions. Reflecting on the significance of the trimer-of-dimer interaction (signaling) domain of WspA^19^, the corresponding region is extremely conserved. This particular region also shares 74.5% sequence similarity to the enteric chemotaxis Tsr signaling domain^68,69^, and we found that the signaling domain in WspA likely extends beyond those reported^19^ to include residues 378-432 (Table S5). Mutations within this signaling domain likely alter the stability of the trimer-of-dimer interactions, resulting in either increased or decreased signaling to WspE^19^. Two additional regions exhibit high conservation (residues 291-300 and 499-508) which we predict to function as the methylation sites (CH3) by WspC (Table S5). Although methylation of WspA has never been experimentally confirmed^62^, high sequence conservation coupled with homology to the chemotaxis system strongly suggests that these sites are likely methylated. Lastly, the chemotaxis Tsr contains a docking site for the methyltransferase CheR (Fig. 1A), which is present as the last five residues of the Tsr C-terminus^70^. Although WspA does not contain the same motif, we observe high conservation of the last 9 amino acids (536-545) at the C-terminus. This region likely represents the docking site for WspC and WspF. Interestingly, there is relatively low sequence conservation in the extracellular sensory domain (Fig. 5). The same pattern holds true across the R-system but the H-system shows much greater levels of conservation (Fig. S3). These results suggest that these two systems may respond to unique extracellular stimuli, which could also apply across the R-system as well given the extent of sequence variations observed.

WspB and WspD are loosely thought to function to relay the signal between WspA and WspE like their chemotaxis CheW homolog^71,72^ (Fig. 1A). However, O’Connor *et al* found that deletions of *wspB* and *wspD* in *P. aeruginosa* manifest discrete outcomes on the subcellular localization of WspA^19^. A Δ*wspD* strain caused WspA to become polarly localized, dramatically reducing WspR phosphorylation. In contrast, a *ΔwspB* strain had little effect on WspA’s naturally lateral presence but resulted in similarly reduced WspR phosphorylation. This indicates that WspB and WspD are not functionally redundant and suggests an essential role for WspD in modulating the subcellular localization of the Wsp system and the intracellular pool of cyclic di-GMP. Our analysis reveals that WspB and WspD are the least conserved among all Wsp proteins and they exhibit unique conservation signatures (Figs. 5 and S4). We identified six regions of WspB where motifs between 8-17 residues in length are highly conserved. WspD similarly has 7 motifs ranging between 5-19 residues. Comparisons of these motifs reveals that they are unique to their respective proteins and have no known or predicted function. Although we were unable to bioinformatically predict functional domains in either protein, their unique conservation signatures described here likely represent sites for interacting with WspA and WspE (Fig. 1A), and potentially with other non-Wsp proteins to modulate subcellular localization. Interestingly, no mutation has ever been reported in the *wspB* gene for the H-system (Fig. 5).

Both the conserved regions and reported mutations in WspC and WspF largely associate with our annotated domains (Fig. 5). WspC is predicted to function as the main activator of the Wsp system and WspF acts as the repressor (Fig. 1A). Consequently, *wspF* mutations from the literature exclusively act to turn on the Wsp system while mutations in *wspC* exclusively turn off the Wsp system (Fig. 5). However, there is a striking pattern here where no mutation has ever been reported for either protein of the H-system. This is very surprising given that WspC and WspF are predicted to function as the main switch of the Wsp system. Collectively, these results suggest that WspC and WspF indeed function to methylate and de-methylate WspA as predicted, but the methylation state of WspA may have reduced influence on the activity of the H-system compared to the R-system.

We observe high conservation in the REC domain (648-702AA) of WspE (Fig. 5) and mutations within specific residues that appear to activate WspE in a WspA-independent manner^18^. The HATPase domain shows high conservation, which is expected as this region is essential for binding to ATP and ultimately phosphorylating WspF and WspR/WspH. We observe high conservation in unannotated regions that flank the HATPase domain that are frequently mutated in the H-system but entirely unaffected in the R-system. Given that WspE interacts with either WspR or WspH, this particular region may be uniquely important for the H-system. It is likely that these WspE activating mutations observed exclusively in the H-system manifest a conformational change that initiates HATPase function in the absence of a stimulus. The region between the HPT and HATPase domains in the chemotaxis CheA constitute the P2 and P3 domains which are responsible for CheY (response regulator) and CheB (methylesterase) docking and CheA dimerization, respectively (Fig. 1A). However, blast assessments reveal no significant similarities between WspE and CheA for these regions, and therefore WspE lacks this annotation. It is possible that the H-system mutations in the 3’ adjacent HATPase region my affect either WspE dimerization or WspH/WspF docking and this may be uniquely important for the H-system.

Previous work compared WspH of *B. cenocepacia* and WspR of *P. aeruginosa* to find ∼70% sequence similarity within the REC domain^43^, suggesting that WspH and WspR are similarly phosphorylated by WspE. However, WspH and WspR differ substantially in their C-termini where WspR contains the diguanylate cyclase GGDEF domain and WspH possesses a histidine kinase speculated to phosphorylate an unknown response regulator^43^. We observe comparably high conservation in our large dataset within the WspH/R REC domain (Fig. 5). As expected, we observe weak conservation of this C-terminus in our WspH/R alignment and greater conservation when H- and R-systems are compared independently (Fig. S3). Interestingly, only the GGDEF region of the WspR C-terminus exhibits high conservation.

### Identification of residues that are uniquely conserved between H- and R-systems

Among the 2,899 consensus amino acid residues of the H- and R-systems, we have identified 43 residues that are likely important for the specialized function of the H-system (see Methods). These 43 residues are highly conserved in both H- and R-systems but also unique to each system (Table 1), and represent non-conservative substitutions that occurred within the H-system^57^. Selective pressures have forced nearly all H-systems to retain these residues, suggesting that they are essential to H-system signaling. Notably, 23 of the 43 identified residues are in the C-terminus of WspH, which does not encode an enzymatic domain like WspR, but is instead a histidine kinase speculated to phosphorylate an unknown response regulator that stimulates biofilm production^43^. Seven residues occur in the REC domain of WspH, which is predicted to interact with and become phosphorylated by WspE. WspE of the H-system has three substitutions, with two occurring in the histidine phosphotransfer (HPT) relay domain and one in an unannotated region of the C-terminus. We predict that these residues are unique to WspE and WspH interactions and WspH phosphorylation. Five substitutions were identified in unannotated regions of WspA, WspB, and WspD, four remaining substitutions were within the methyltransferase domain of WspC, and none were detected in WspF. These uniquely conserved substitutions likely differentiate the signaling mechanism of the H-system from the R-system, which remains entirely unexplored.

**Table 1.**
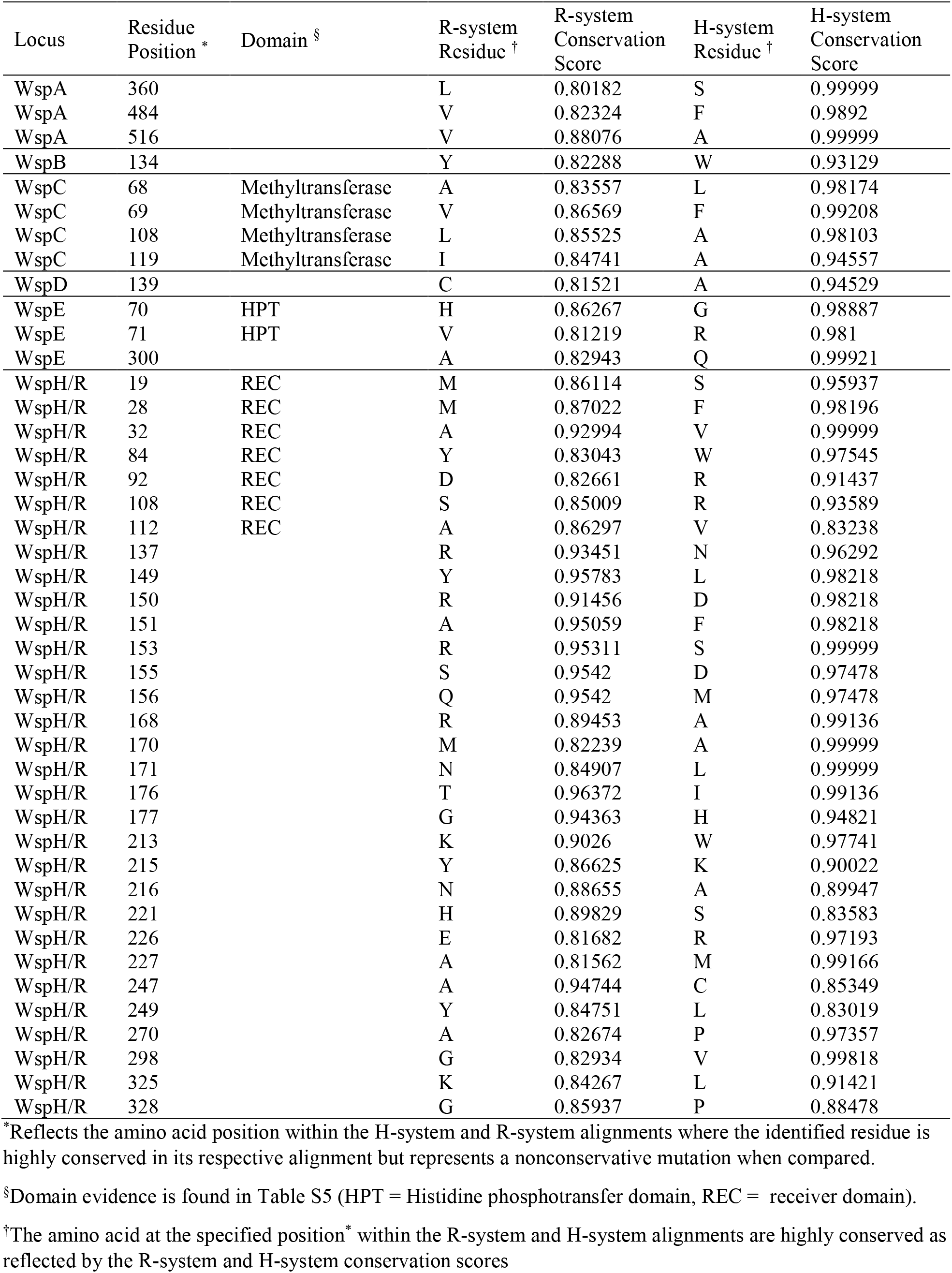
Assessment of unique conservation between the H- and R-systems identifies residues important for the specialized function of the H-system.

| Locus | Residue Position * | Domain § | R-system Residue † | R-system Conservation Score | H-system Residue † | H-system Conservation Score |
| --- | --- | --- | --- | --- | --- | --- |
| WspA | 360 |  | L | 0.80182 | S | 0.99999 |
| WspA | 484 |  | V | 0.82324 | F | 0.9892 |
| WspA | 516 |  | V | 0.88076 | A | 0.99999 |
| WspB | 134 |  | Y | 0.82288 | W | 0.93129 |
| WspC | 68 | Methyltransferase | A | 0.83557 | L | 0.98174 |
| WspC | 69 | Methyltransferase | V | 0.86569 | F | 0.99208 |
| WspC | 108 | Methyltransferase | L | 0.85525 | A | 0.98103 |
| WspC | 119 | Methyltransferase | I | 0.84741 | A | 0.94557 |
| WspD | 139 |  | C | 0.81521 | A | 0.94529 |
| WspE | 70 | HPT | H | 0.86267 | G | 0.98887 |
| WspE | 71 | HPT | V | 0.81219 | R | 0.981 |
| WspE | 300 |  | A | 0.82943 | Q | 0.99921 |
| WspH/R | 19 | REC | M | 0.86114 | S | 0.95937 |
| WspH/R | 28 | REC | M | 0.87022 | F | 0.98196 |
| WspH/R | 32 | REC | A | 0.92994 | V | 0.99999 |
| WspH/R | 84 | REC | Y | 0.83043 | W | 0.97545 |
| WspH/R | 92 | REC | D | 0.82661 | R | 0.91437 |
| WspH/R | 108 | REC | S | 0.85009 | R | 0.93589 |
| WspH/R | 112 | REC | A | 0.86297 | V | 0.83238 |
| WspH/R | 137 |  | R | 0.93451 | N | 0.96292 |
| WspH/R | 149 |  | Y | 0.95783 | L | 0.98218 |
| WspH/R | 150 |  | R | 0.91456 | D | 0.98218 |
| WspH/R | 151 |  | A | 0.95059 | F | 0.98218 |
| WspH/R | 153 |  | R | 0.95311 | S | 0.99999 |
| WspH/R | 155 |  | S | 0.9542 | D | 0.97478 |
| WspH/R | 156 |  | Q | 0.9542 | M | 0.97478 |
| WspH/R | 168 |  | R | 0.89453 | A | 0.99136 |
| WspH/R | 170 |  | M | 0.82239 | A | 0.99999 |
| WspH/R | 171 |  | N | 0.84907 | L | 0.99999 |
| WspH/R | 176 |  | T | 0.96372 | I | 0.99136 |
| WspH/R | 177 |  | G | 0.94363 | H | 0.94821 |
| WspH/R | 213 |  | K | 0.9026 | W | 0.97741 |
| WspH/R | 215 |  | Y | 0.86625 | K | 0.90022 |
| WspH/R | 216 |  | N | 0.88655 | A | 0.89947 |
| WspH/R | 221 |  | H | 0.89829 | S | 0.83583 |
| WspH/R | 226 |  | E | 0.81682 | R | 0.97193 |
| WspH/R | 227 |  | A | 0.81562 | M | 0.99166 |
| WspH/R | 247 |  | A | 0.94744 | C | 0.85349 |
| WspH/R | 249 |  | Y | 0.84751 | L | 0.83019 |
| WspH/R | 270 |  | A | 0.82674 | P | 0.97357 |
| WspH/R | 298 |  | G | 0.82934 | V | 0.99818 |
| WspH/R | 325 |  | K | 0.84267 | L | 0.91421 |
| WspH/R | 328 |  | G | 0.85937 | P | 0.88478 |

\*Reflects the amino acid position within the H-system and R-system alignments where the identified residue is highly conserved in its respective alignment but represents a nonconservative mutation when compared.
§Domain evidence is found in Table S5 (HPT = Histidine phosphotransfer domain, REC = receiver domain).
†The amino acid at the specified position\* within the R-system and H-system alignments are highly conserved as reflected by the R-system and H-system conservation scores

## Conclusion

This study presents the first broad phylogenetic analysis of the Wsp signaling system to date. This research expands our current understanding of the Wsp signaling system which had been thus far restricted to four *Pseudomonas* spp. and more recently to *B. cenocepacia* HI2424. Ironically, we have found that *Pseudomonas* spp. are the only γ-proteobacteria to possess the Wsp system, which appears to have originated in the β-proteobacteria with strong indications of multiple horizontal gene transfer events. There are clear genetic and biochemical evidence ^19,27,28,32–34,37,39,45–47^ to suggest that the core signaling components of the Wsp system in *Pseudomonas* mechanistically function much like those of the enteric chemotaxis system. Our sequence conservation and mutation analysis extends our current understanding of Wsp to include more evolutionary divergent organisms and reveals strong potential for mechanistic diversity among individual Wsp systems.

All Wsp systems appear to utilize either WspH or WspR exclusively as the terminal output of the signaling cascade, and all organisms possess only one H- or R-system. An earlier study discovered the H-system in *B. cenocepacia* HI2424 to show that WspH lacks the diguanylate cyclase GGDEF domain found in WspR and speculated WspH likely activates a unique diguanylate cyclase^43^. This was based on the observation that various mutations in *wsp* genes caused the hallmark wrinkly colony morphology and increased biofilm formation associated with elevated cyclic di-GMP production in diverse bacterial species. However, our study clearly demonstrates that these *wsp* mutations in *B. cenocepacia* HI2424 do not alter cyclic di-GMP production in contrast to comparable *wsp* mutations in *P. fluorescens* Pf0-1. Despite the striking difference in the functional output of H- and R-systems, they share high levels of sequence conservation within and beyond key functional domains that overlap with the enteric chemotaxis system. One major difference we observed was the complete absence of reported mutations in *wspC* and *wspF* genes of the H-system in contrast to those of the R-system. We also identified 43 highly conserved amino acid residues across all Wsp proteins that are uniquely modified in the H-system. These specific residues likely differentiate mechanistic variations between the H- and R-systems, and we suspect that the methylation state of WspA exerts a relatively reduced role in the signaling cascade of the H-system compared to the R-system.

The Wsp proteins of the R-system exhibit greater overall sequence variation compared to those of the H-system. This is not surprising since the H-system is phylogenetically restricted to *Burkholderia* spp. and the R-system is much more divergent. However, nearly all of the annotated functional domains are highly conserved across the R-system with the exception being the extracellular sensory domain of WspA. The external stimulus of the Wsp system long remained a mystery until a recent study demonstrated surface-contact to be the main stimulus in *P. aeruginosa*^38^. The relatively large sequence variation observed within this extracellular sensory domain indicates strong potential for independent adaptations to diverse external stimuli. We found that WspB and WspD proteins exhibit the least amount of sequence conservation, yet we did not observe any instance of a *wsp* operon lacking either of these proteins, strongly indicating that they are both functionally important. In contrast to the enteric chemotaxis system which utilizes CheW to physically bridge the methylation and phosphorylation signaling modules, the Wsp system is thought to utilize both WspB and WspD in an analogous manner. However, there is clear evidence in *P. aeruginosa* that WspB and WspD are not functionally redundant^19^ and we observed distinct sequence conservation patterns between these two proteins. We also found that all mutations reported in WspB and WspD proteins act to deactivate the R-system while mutations in WspD of the H-system exclusively act to turn it on. Furthermore, no mutations have been reported for WspB of the H-system. Given the tremendous influence of WspD to *P. aeruginosa*’s Wsp signaling cascade^19^, these so called accessory proteins likely play both functionally important and diverse roles among individual Wsp systems.

Mutations in *wsp* genes are frequently identified in clinical specimens and are associated with increased biofilm formation^22,24,25,73–78^, whether or not this is achieved through elevated production of cyclic di-GMP. Genetically deregulating the Wsp system appears to be a significant but transient adaptive strategy in a dynamic and overcrowded ecosystem. Our study shows that leaning on functional analogies to the enteric chemotaxis system serves as both a valid and critical foundation for future studies to understand the functional blueprint of diverse Wsp systems in greater detail. Key focus should be placed on elucidating the function of WspH and its interacting partners, disentangling the functional uniqueness of WspB and WspD, and identifying extracellular stimuli for WspA in diverse organisms.

## Supplementary Information

Figs S1-S4 and Tables S1-S5

Data S1. MultiGeneBlast metadata for the *P. fluorescens* Pf0-1 *wsp* operon.

Data S2. MultiGeneBlast metadata for the *B. cenocepacia* HI2424 *wsp* gene cluster.

File S1. Python script executed on Data S1-S2.

File S2. R script executed by File S1 to download RefSeq genome assemblies.

File S3. R script executed by File S1 to parse File S2 output for Bac120 set creation.

File S4. R script executed by File S1 to parse File S3 output and finalize Bac120 dataset.

## Acknowledgements

We thank M. Bentley for developing the initial codes for phylogenetic analyses. This study was funded by the Charles Henry Leach II Fund (W.K.) and the National Institute of General Medical Sciences of the NIH 1R15GM132856 (W.K.).

## Author Contributions

C.K and W.K. designed the study, analyzed data, and wrote the manuscript; C.K. performed experiments; and E.M. and V.C. provided strains and advice.

## Supplementary Figures and Tables

**Figure S1.**
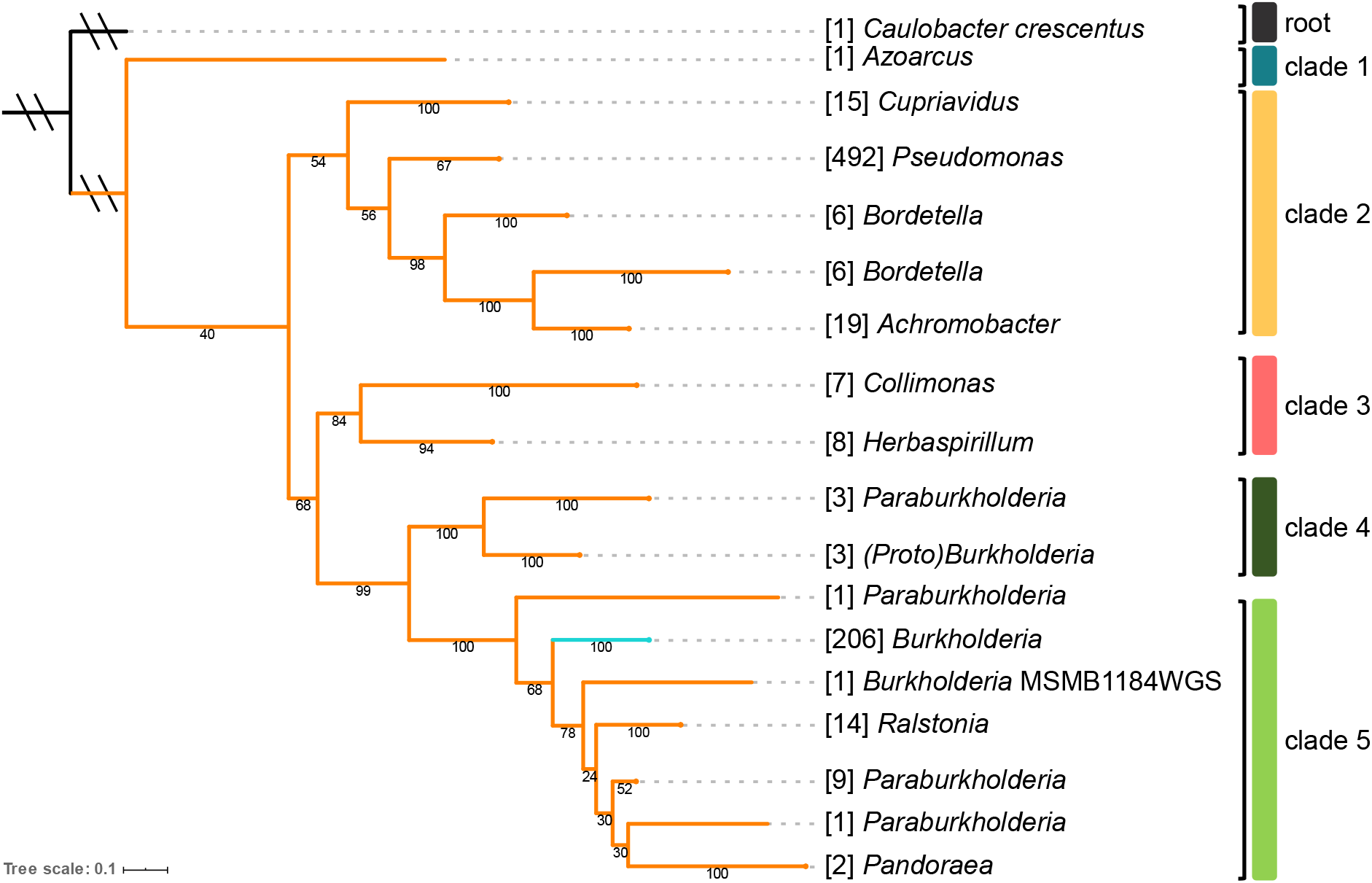
Phylogenetic analysis of the core Wsp proteins indicates that the R-system predates the H-system. The H-system (teal) and R-system (orange) gene tree was constructed using the amino acid sequences of the core Wsp proteins (WspA, WspB, WspC, WspD, WspE, and WspF). The phylogeny is rooted to Wsp homologs in *Caulobacter crescentus* reported in Table S2. The Wsp system diverges into five distinct clades. Each clade is assigned a unique color and the same color scheme is applied to Figs. 3 and 4 to denote the respective clades. The R-system precedes the H-system and all 206 H-systems form a single branch. The values within parentheses indicate the number of species/strains within each branch and those under each branch represent the bootstrap support values (out of 100).

**Figure S2.**
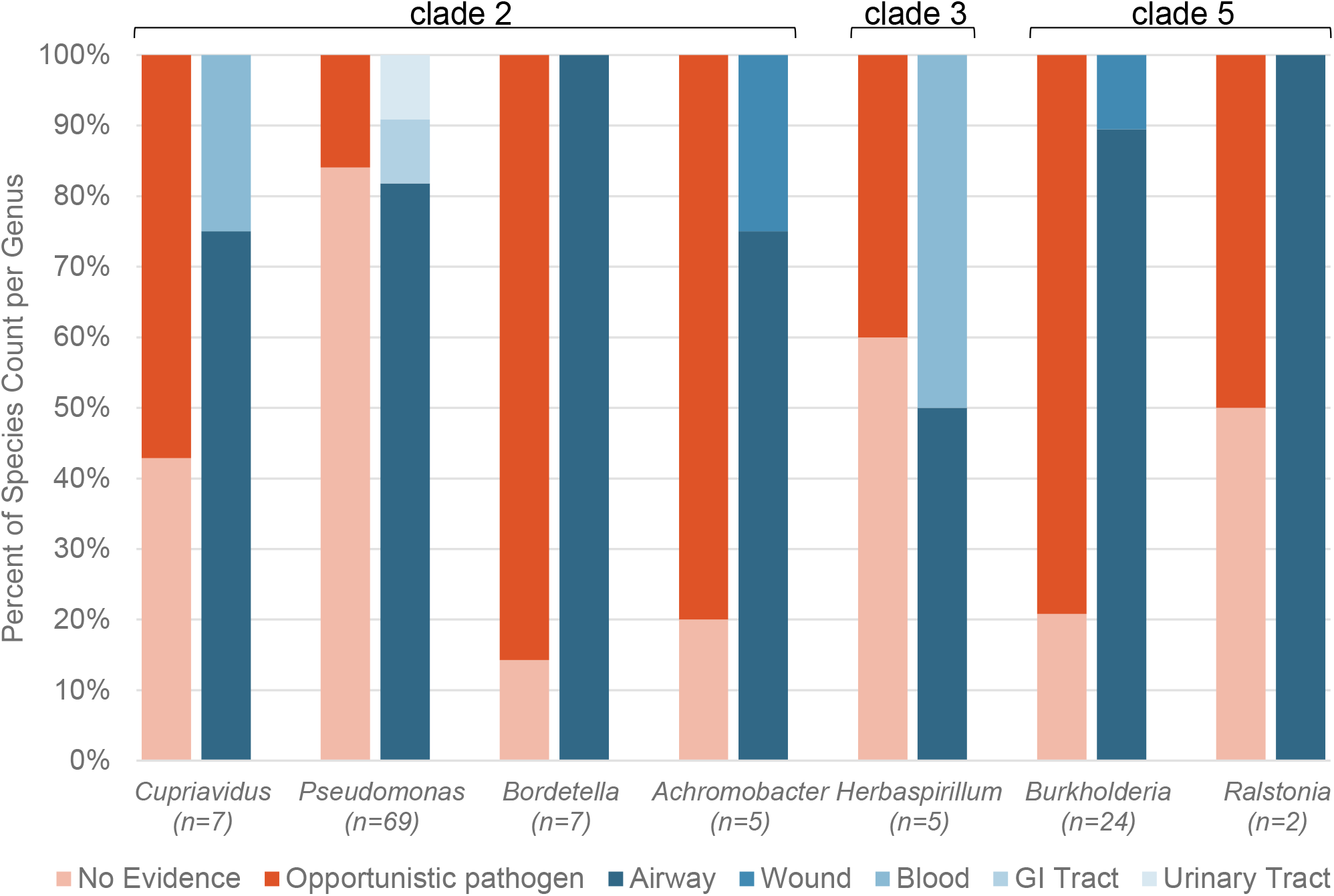
Opportunistic pathogens with a Wsp system are frequently associated with the respiratory environment. Organisms represented in the Wsp dataset were classified as either opportunistic pathogens (red) or no evidence of pathogenicity (rose) based on a literature search at the species level. Species identified as opportunistic pathogens were then sub-categorized by common site of isolation (blue). Counts of unique species for each genus are reported on the x-axis. Genera without any indication of pathogenicity were excluded from this figure but the full analysis with references is summarized in Table S3.

**Figure S3.**
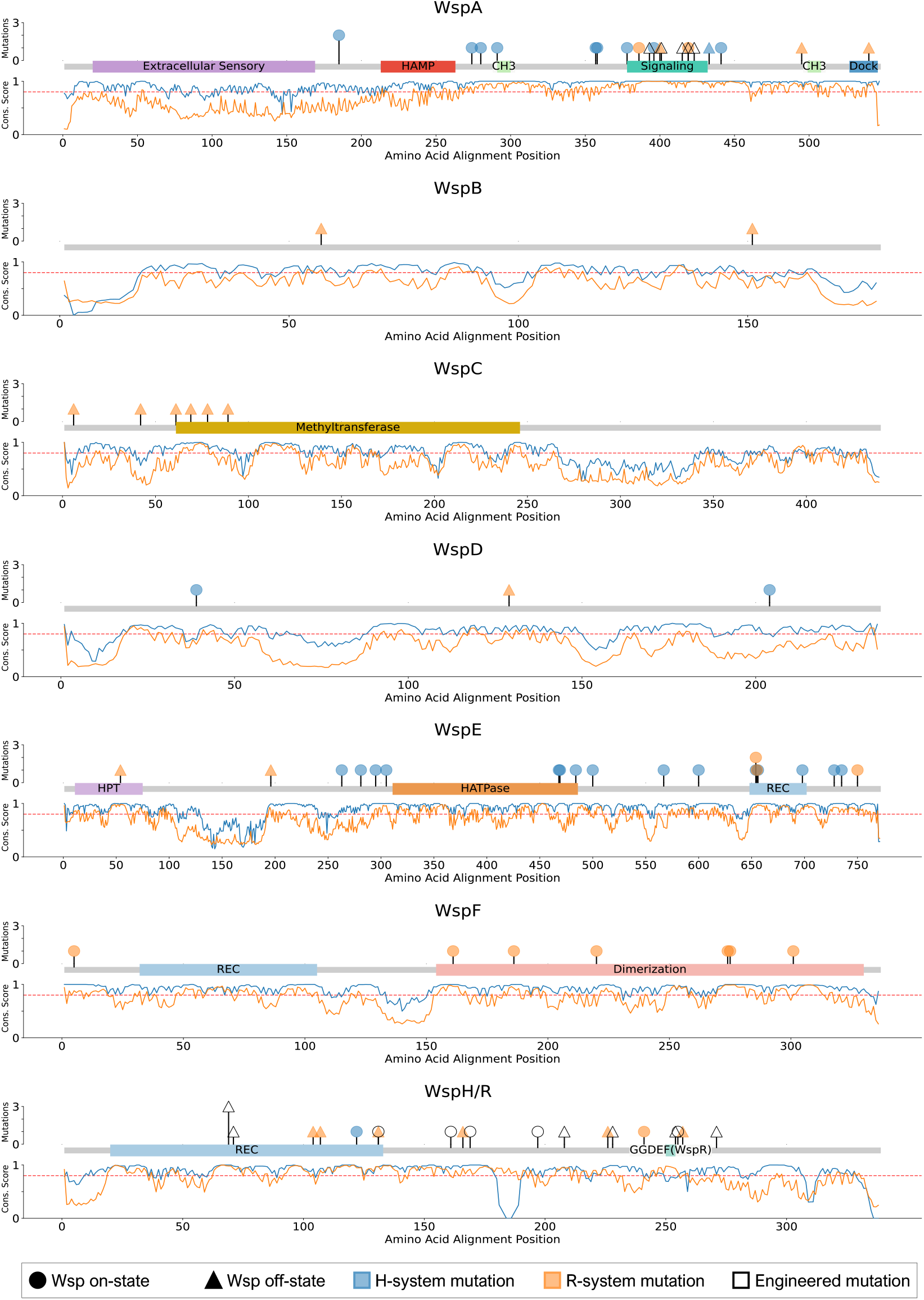
Discrete conservation of Wsp proteins within H- or R-systems. Amino acid sequences used to generate Figure 5 were divided into an H-system plot (blue) or an R-system plot (orange). Annotation data is derived from NCBI CDD (conserved domain database), Prosite, or Che homology as indicated in Table S5. Reported naturally occurring missense mutations from the literature in the H-system are shown in blue and those in the R-system are shown in orange. Engineered missense mutations reported in the literature are indicated in white. Mutations that turn on the respective Wsp system are indicated as circles while those that turn off the system are indicated as triangles. The y-axis represents the Shannon Entropy evaluation for each protein alignment where weighted values near 1 indicate high sequence conservation and values near zero indicate weak sequence conservation. The horizontal dashed line indicates where the weighted Shannon Entropy metric equals 0.8, denoting residues of greater functional or structural importance.

**Figure S4.**
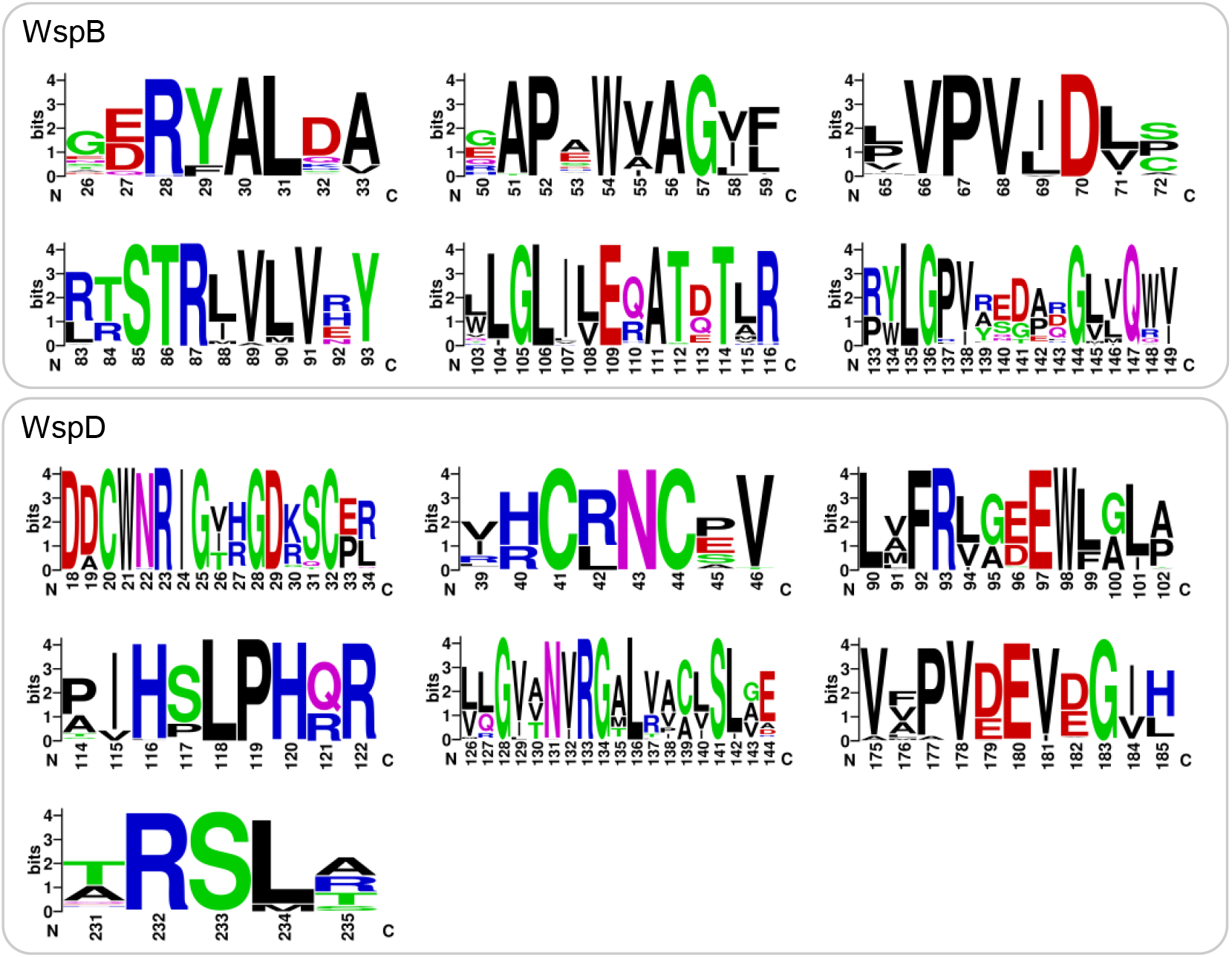
Sequence logos of conserved regions in WspB and WspD. Individual domains of high conservation between WspB and WspD show little to no similarity. Sequence logos were generated to represent the highly conserved regions in Figure 5 for WspB and WspD. Islands of high conservation were identified by the 0.8 Shannon entropy metric. Two residues immediately flanking the conserved region are included in the Sequence logos to ensure the entire conserved region is depicted. The values indicated on the x-axis denote the relative amino acid position within the coding sequence. No overlapping signatures were found between the WspB and WspD, which is surprising given their proposed similar function.

**Table S1.** Sequence conservation assessment of the Wsp signal transduction system and the enteric chemotaxis Che system.

| Protein Identifier | *E. coli* | *P. fluorescens* | *B. cenocepacia* | Average Conservation Score† |  |
| --- | --- | --- | --- | --- | --- |
|  |  |  |  | Comparison of all 3 systems | Comparison of the H- and R-systems |
| Tsr/WspA | NP_415938 | ABA72795 | ABK10523 | 0.334 | 0.521 |
| CheW/WspB | NP_416401 | ABA72796 | ABK10524 | 0.285 | 0.383 |
| CheR/WspC | NP_416938 | ABA72797 | ABK10525 | 0.328 | 0.330 |
| CheW/WspD | NP_416401 | ABA72798 | ABK10527 | 0.278 | 0.319 |
| CheA/WspE | NP_416402 | ABA72799 | ABK10528 | 0.342 | 0.429 |
| CheB/WspF | NP_416397 | ABA72800 | ABK10529 | 0.412 | 0.584 |
| CheY/WspR/WspH | NP_416396 | ABA72801 | ABK10522 | 0.328 | 0.268 |

† Conservation scores assessed through Shannon entropy analyses (see Methods). Score ranges from 0 to 1 with 0 indicating no conservation and 1 indicating complete conservation. The score was determined for each residue in the alignment. The average conservation score of the protein alignment is reported for each system comparison.

**Table S2.** Establishing *C. crescentus* Wsp homologs as the root for the Wsp phylogenetic analysis.

| *C. crescentus* | NCBI Accession | Homolog | Query † | Query Cover | E-value § | % Identity |
| --- | --- | --- | --- | --- | --- | --- |
| CP001340.1 | ACL96924 | WspA | Pfl01_1052 | 60% | 3.00E-29 | 29.31% |
| CP001340.1 | ACL96585 | WspB | Pfl01_1053 | 72% | 4.00E-08 | 27.82% |
| CP001340.1 | ACL93911 | WspC | Pfl01_1054 | 44% | 8.00E-25 | 32.66% |
| CP001340.1 | ACL94859 | WspD | Pfl01_1055 | 19% | 1.00E-04 | 31.82% |
| CP001340.1 | ACL94095 | WspE | Pfl01_1056 | 78% | 7.00E-68 | 30.36% |
| CP001340.1 | ACL93912 | WspF | Pfl01_1057 | 99% | 1.00E-49 | 34.01% |

† *P. fluorescens* Pf0-1 Wsp sequences were used as a query against *C. crescentus* CP001340.1 to identify respective homologs for rooting the Wsp phylogeny.
§ Identified proteins scored low E-values, indicating that they are acceptable homologs for rooting the Wsp phylogeny.

**Table S3.** A literature review of individual species within the Wsp dataset identifies opportunistic pathogens.

| Species * | Count ** | Human Pathogen Type † | Site of Infection †† | Reference § | Species * | Count ** | Human Pathogen Type † |
| --- | --- | --- | --- | --- | --- | --- | --- |
| *Achromobacter denitrificans* | 3 | Opportunistic pathogen | Airway | Awadh *et al.* (2017) | *Paraburkholderia caribensis* | 4 | No Evidence |
| *Achromobacter insolitus* | 4 | Opportunistic pathogen | Wound | Coenye *et al.* (2003) | *Paraburkholderia fungorum* | 1 | No Evidence |
| *Achromobacter spanius* | 4 | Opportunistic pathogen | Airway | Edwards *et al.* (2017) | *Paraburkholderia hospita* | 2 | No Evidence |
| *Achromobacter xylosoxidans* | 5 | Opportunistic pathogen | Airway | Awadh *et al.* (2017) | *Paraburkholderia phymatum* | 1 | No Evidence |
| *Bordetella avium* | 3 | Opportunistic pathogen | Airway | Harrington *et al.* (2009) | *Paraburkholderia phytofirmans* | 1 | No Evidence |
| *Bordetella bronchialis* | 2 | Opportunistic pathogen | Airway | Vandamme *et al.* (2015) | *Paraburkholderia sp.* | 2 | No Evidence |
| *Bordetella flabialis* | 1 | Opportunistic pathogen | Airway | Linz *et al.* (2019) | *Paraburkholderia terrae* | 1 | No Evidence |
| *Bordetella genomosp.* | 1 | Opportunistic pathogen | Airway | Spilker *et al.* (2017) | *Pseudomonas alkylphenolica* | 1 | No Evidence |
| *Bordetella hinzii* | 3 | Opportunistic pathogen | Airway | Cookson *et al.* (1994) | *Pseudomonas amygdali* | 2 | No Evidence |
| *Bordetella sp.* | 1 | Opportunistic pathogen | Airway | Weigand *et al.* (2017) | *Pseudomonas antarctica* | 2 | No Evidence |
| *Burkholderia ambifaria* | 1 | Opportunistic pathogen | Airway | Sousa *et al.* (2011) | *Pseudomonas arsenicoxydans* | 1 | No Evidence |
| *Burkholderia cenocepacia* | 18 | Opportunistic pathogen | Airway | García-Romero *et al.* (2020) | *Pseudomonas asplenii* | 1 | No Evidence |
| *Burkholderia cepacia* | 8 | Opportunistic pathogen | Airway | McClean *et al.* (2009) | *Pseudomonas avellanae* | 1 | No Evidence |
| *Burkholderia contaminans* | 2 | Opportunistic pathogen | Airway | Sousa *et al.* (2011) | *Pseudomonas azotoformans* | 3 | No Evidence |
| *Burkholderia diffusa* | 1 | Opportunistic pathogen | Airway | Sousa *et al.* (2011) | *Pseudomonas brassicaearum* | 5 | No Evidence |
| *Burkholderia dolosa* | 2 | Opportunistic pathogen | Airway | Sousa *et al.* (2011) | *Pseudomonas brenneri* | 1 | No Evidence |
| *Burkholderia glumae* | 2 | Opportunistic pathogen | Airway | Wineberg *et al.* (2007) | *Pseudomonas cerasi* | 2 | No Evidence |
| *Burkholderia lata* | 3 | Opportunistic pathogen | Airway | Leong *et al.* (2018) | *Pseudomonas chlororaphis* | 46 | No Evidence |
| *Burkholderia mallei* | 21 | Opportunistic pathogen | Airway | Saikh *et al.* (2017) | *Pseudomonas citronellolis* | 2 | No Evidence |
| *Burkholderia metallica* | 1 | Opportunistic pathogen | Airway | Sousa *et al.* (2011) | *Pseudomonas corrugata* | 2 | No Evidence |
| *Burkholderia multivorans* | 8 | Opportunistic pathogen | Airway | Rhodes *et al.* (2016) | *Pseudomonas cremoricolorata* | 1 | No Evidence |
| *Burkholderia oklahomensis* | 2 | Opportunistic pathogen | Wound | Glass *et al.* (2006) | *Pseudomonas entomophila* | 3 | No Evidence |
| *Burkholderia pseudomallei* | 92 | Opportunistic pathogen | Airway | Broek *et al.* (2017) | *Pseudomonas extremaustralis* | 1 | No Evidence |
| *Burkholderia pyrocinia* | 2 | Opportunistic pathogen | Airway | Sousa *et al.* (2011) | *Pseudomonas extremorientalis* | 1 | No Evidence |
| *Burkholderia seminalis* | 1 | Opportunistic pathogen | Airway | Sousa *et al.* (2011) | *Pseudomonas fluorescens* | 22 | No Evidence |
| *Burkholderia stabilis* | 3 | Opportunistic pathogen | Airway | Sousa *et al.* (2011) | *Pseudomonas fragi* | 4 | No Evidence |
| *Burkholderia thailandensis* | 16 | Opportunistic pathogen | Wound | Gee *et al.* (2018) | *Pseudomonas frederiksbergensis* | 3 | No Evidence |
| *Burkholderia ubonensis* | 1 | Opportunistic pathogen | Airway | Sousa *et al.* (2011) | *Pseudomonas furukawaii* | 1 | No Evidence |
| *Burkholderia vietnamiensis* | 5 | Opportunistic pathogen | Airway | Jassem *et al.* (2011) | *Pseudomonas fuscovaginae* | 1 | No Evidence |
| *Cupriavidus metallidurans* | 2 | Opportunistic pathogen | Blood | Monsieurs *et al.* (2013) | *Pseudomonas granadensis* | 1 | No Evidence |
| *Cupriavidus necator* | 1 | Opportunistic pathogen | Airway | Zhang *et al.* (2016) | *Pseudomonas knackmussii* | 1 | No Evidence |
| *Cupriavidus pauculus* | 1 | Opportunistic pathogen | Airway | Almasy *et al.* (2016) | *Pseudomonas koreensis* | 3 | No Evidence |
| *Cupriavidus taiwanensis* | 8 | Opportunistic pathogen | Airway | Chen *et al.* (2001) | *Pseudomonas kribbensis* | 1 | No Evidence |
| *Herbaspirillum huttiense* | 1 | Opportunistic pathogen | Blood | Hernández *et al.* (2019) | *Pseudomonas libanensis* | 1 | No Evidence |
| *Herbaspirillum seropedicae* | 3 | Opportunistic pathogen | Airway | Dhital *et al.* 2020 | *Pseudomonas lini* | 1 | No Evidence |
| *Pseudomonas aeruginosa* | 181 | Opportunistic pathogen | Airway | Fujitani *et al.* (2011) | *Pseudomonas lurida* | 2 | No Evidence |
| *Pseudomonas fulva* | 2 | Opportunistic pathogen | Airway | Shariff *et al.* (2017) | *Pseudomonas mandelii* | 2 | No Evidence |
| *Pseudomonas mendocina* | 4 | Opportunistic pathogen | Airway | Ioannou *et al.* (2020) | *Pseudomonas mediterranea* | 1 | No Evidence |
| *Pseudomonas monteilii* | 5 | Opportunistic pathogen | Airway | Shariff *et al.* (2017) | *Pseudomonas moraviensis* | 1 | No Evidence |
| *Pseudomonas mosselii* | 1 | Opportunistic pathogen | Airway | Shariff *et al.* (2017) | *Pseudomonas mucidolens* | 2 | No Evidence |
| *Pseudomonas plecoglossicida* | 2 | Opportunistic pathogen | Airway | Shariff *et al.* (2017) | *Pseudomonas orientalis* | 6 | No Evidence |
| *Pseudomonas putida* | 27 | Opportunistic pathogen | Airway | Shariff *et al.* (2017) | *Pseudomonas palleroniana* | 1 | No Evidence |
| *Pseudomonas rhodesiae* | 1 | Opportunistic pathogen | Airway | Sawa *et al.* (2016) | *Pseudomonas parafulva* | 3 | No Evidence |
| *Pseudomonas simiae* | 3 | Opportunistic pathogen | Airway | Vela *et al.* (2006) | *Pseudomonas poae* | 2 | No Evidence |
| *Pseudomonas sp.* | 1 | Opportunistic pathogen | Urinary Tract | McGann *et al.* (2015) | *Pseudomonas prosekii* | 1 | No Evidence |
| *Pseudomonas sp.* | 1 | Opportunistic pathogen | GI Tract | Campos *et al.* (2019) | *Pseudomonas protegens* | 6 | No Evidence |
| *Ralstonia insidiosa* | 2 | Opportunistic pathogen | Airway | Ryan *et al.* (2014) | *Pseudomonas psychrophila* | 1 | No Evidence |
| *Achromobacter sp.* | 3 | No Evidence |  |  | *Pseudomonas reinekei* | 1 | No Evidence |
| *Azoarcus sp.* | 1 | No Evidence |  |  | *Pseudomonas resinovorans* | 1 | No Evidence |
| *Bordetella sp.* | 1 | No Evidence |  |  | *Pseudomonas savastanoi* | 2 | No Evidence |
| *Burkholderia insecticola* | 1 | No Evidence |  |  | *Pseudomonas silesiensis* | 1 | No Evidence |
| *Burkholderia plantarii* | 2 | No Evidence |  |  | *Pseudomonas soli* | 1 | No Evidence |
| *Burkholderia pseudomultivorans* | 1 | No Evidence |  |  | *Pseudomonas sp.* | 64 | No Evidence |
| *Burkholderia sp.* | 17 | No Evidence |  |  | *Pseudomonas synxantha* | 9 | No Evidence |
| *Burkholderia territorii* | 1 | No Evidence |  |  | *Pseudomonas syringae* | 30 | No Evidence |
| *Collimonas arenae* | 3 | No Evidence |  |  | *Pseudomonas taetrolens* | 2 | No Evidence |
| *Collimonas fungivorans* | 2 | No Evidence |  |  | *Pseudomonas thivervalensis* | 1 | No Evidence |
| *Collimonas pratensis* | 2 | No Evidence |  |  | *Pseudomonas tolaasii* | 1 | No Evidence |
| *Cupriavidus basilensis* | 1 | No Evidence |  |  | *Pseudomonas trivialis* | 2 | No Evidence |
| *Cupriavidus oxalaticus* | 1 | No Evidence |  |  | *Pseudomonas umsongensis* | 1 | No Evidence |
| *Cupriavidus pinatubonensis* | 1 | No Evidence |  |  | *Pseudomonas vancouverensis* | 1 | No Evidence |
| *Herbaspirillum hiltneri* | 1 | No Evidence |  |  | *Pseudomonas veronii* | 1 | No Evidence |
| *Herbaspirillum rubrisubalbicans* | 2 | No Evidence |  |  | *Pseudomonas versuta* | 1 | No Evidence |
| *Herbaspirillum sp.* | 1 | No Evidence |  |  | *Pseudomonas viridiflava* | 1 | No Evidence |
| *Pandoraea thiooxydans* | 2 | No Evidence |  |  | *Pseudomonas yamanorum* | 2 | No Evidence |
| *Paraburkholderia aromaticivorans* | 1 | No Evidence |  |  | *Ralstonia solanacearum* | 12 | No Evidence |

\* Species level classification reported on NCBI for the GenBank sequence identified in our analyses.
\*\* The count of strains or isolates in the dataset that fall within the reported species level classification (values sum to 794).
† Species level classification as an opportunistic pathogen or no evidence of pathogenicity based on clinical reports.
†† Location of infection in the clinical reports for strains identified as pathogens or opportunists.
§ Clinical reports, reviews of clinical findings, or NCBI genome submissions that reported the identified species (\*) within a human host during an infection.

**Table S4.**
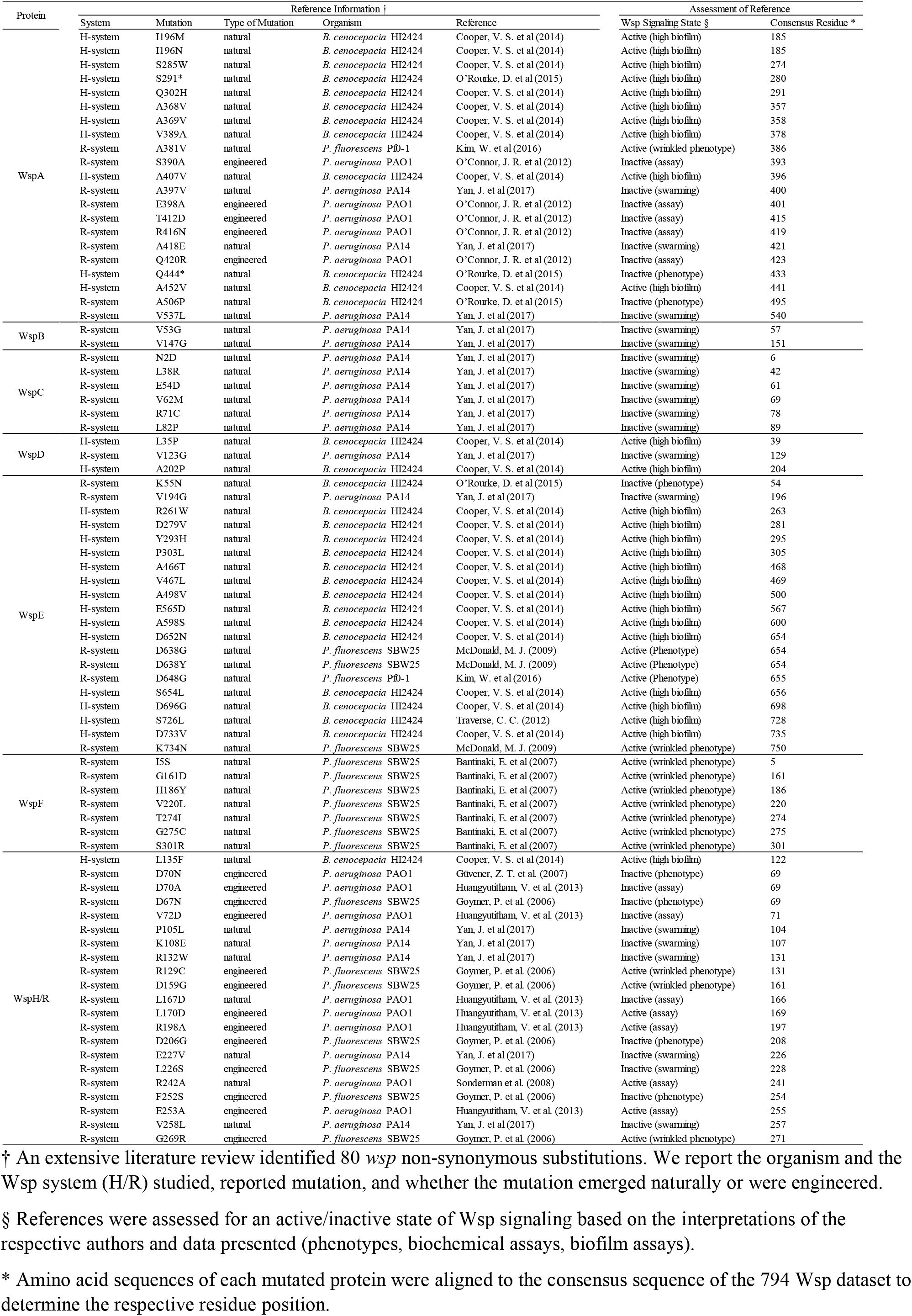
Previously reported non-synonymous substitutions in Wsp systems and associated phenotypes.

**Table S5.**
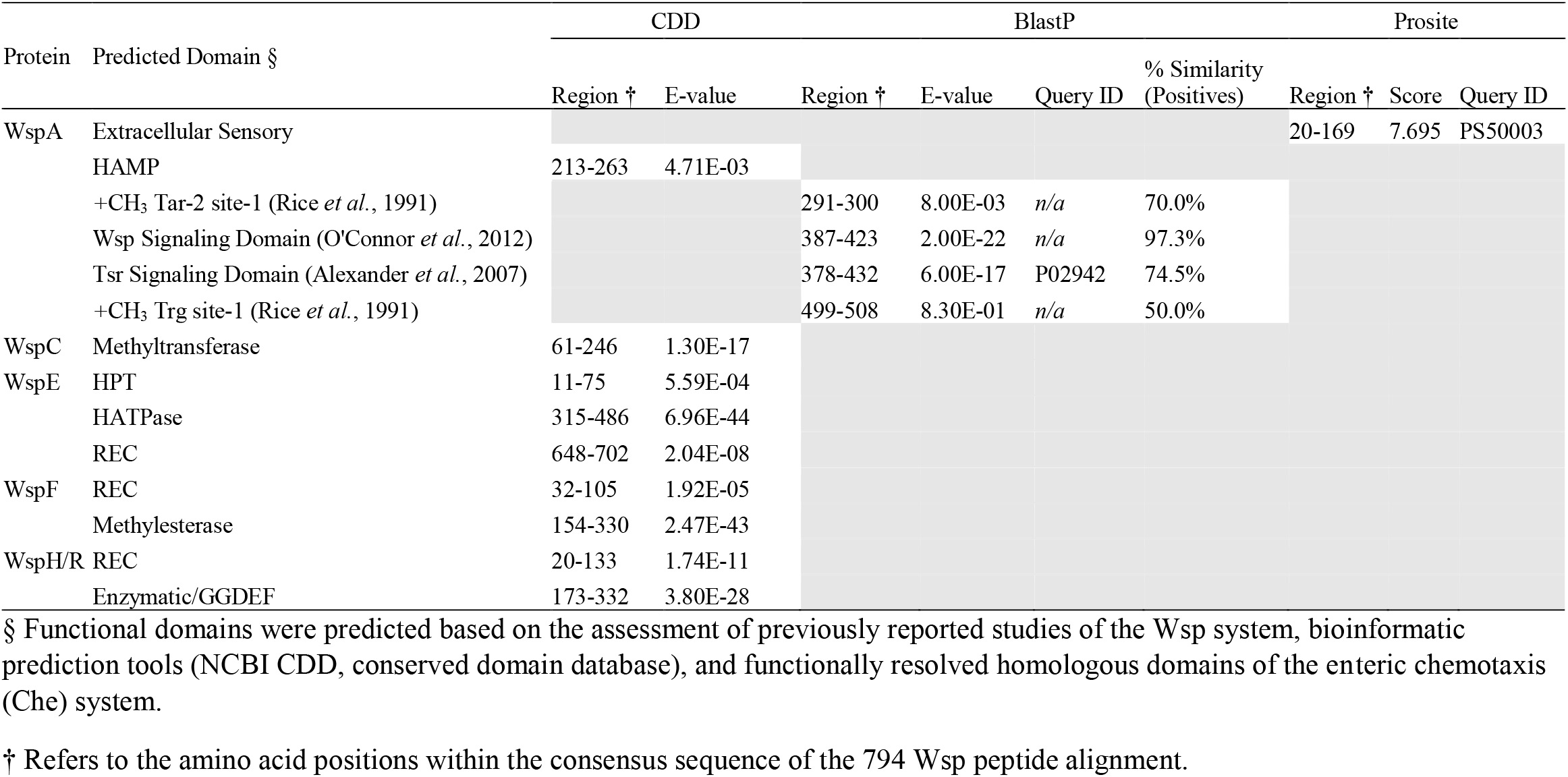
Annotation of functional domains within the consensus sequence of the Wsp signaling system.

